# Site-Specific Incorporation of Citrulline into Proteins in Mammalian Cells

**DOI:** 10.1101/2020.06.06.137885

**Authors:** Santanu Mondal, Shu Wang, Yunan Zheng, Sudeshna Sen, Abhishek Chatterjee, Paul R. Thompson

**Affiliations:** Department of Biochemistry and Molecular Pharmacology, UMass Medical School, 364 Plantation Street, Worcester, MA 01605, USA; Department of Chemistry, Boston College, Chestnut Hill, Massachusetts 02467, USA

## Abstract

Citrullination is a post-translational modification (PTM) of arginine that is crucial for several physiological processes, including gene regulation and neutrophil extracellular trap formation. Despite recent advances, studies of protein citrullination remain challenging due to the difficulty of accessing proteins homogeneously citrullinated at a specific site. Herein, we report a novel technology that enables the site-specific incorporation of citrulline (Cit) into proteins in mammalian cells. This approach exploits an *E. coli*-derived engineered leucyl tRNA synthetase-tRNA pair that incorporates a photocaged-citrulline (**SM60**) into proteins in response to a nonsense codon. Subsequently, **SM60** is readily converted to Cit with light *in vitro* and in living cells. To demonstrate the utility of the method, we biochemically characterized the effect of incorporating Cit at two known autocitrullination sites in Protein Arginine Deiminase 4 (PAD4, R372 and R374) and showed that the R372Cit and R374Cit mutants are 181- and 9-fold less active than the wild-type enzyme. This powerful technology possesses immense potential to decipher the biology of citrullination.

## Introduction

Citrullination is a post-translational modification (PTM) that involves the hydrolysis of the positively-charged guanidium group on arginine to generate a neutral urea (Figure 1a).^1^ Citrullination plays crucial roles in many physiological processes, including the epigenetic regulation of gene transcription, Neutrophil Extracellular Trap (NET)-formation or NETosis, and maintaining pluripotency.^1–7^ Citrullination is catalyzed by the Protein Arginine Deiminases (PADs) (Figure 1a), a group of four catalytically active cysteine hydrolases (PAD1-4).^8^ PADs are Ca^2+^-dependent enzymes and the presence of calcium increases PAD activity by >10,000-fold. Calcium-binding leads to dramatic conformational rearrangements, particularly of the nucleophilic cysteine (C645 in PAD1, 4; C647 in PAD2; C646 in PAD3) to form a catalytically competent active site.^9^

**Figure 1.**
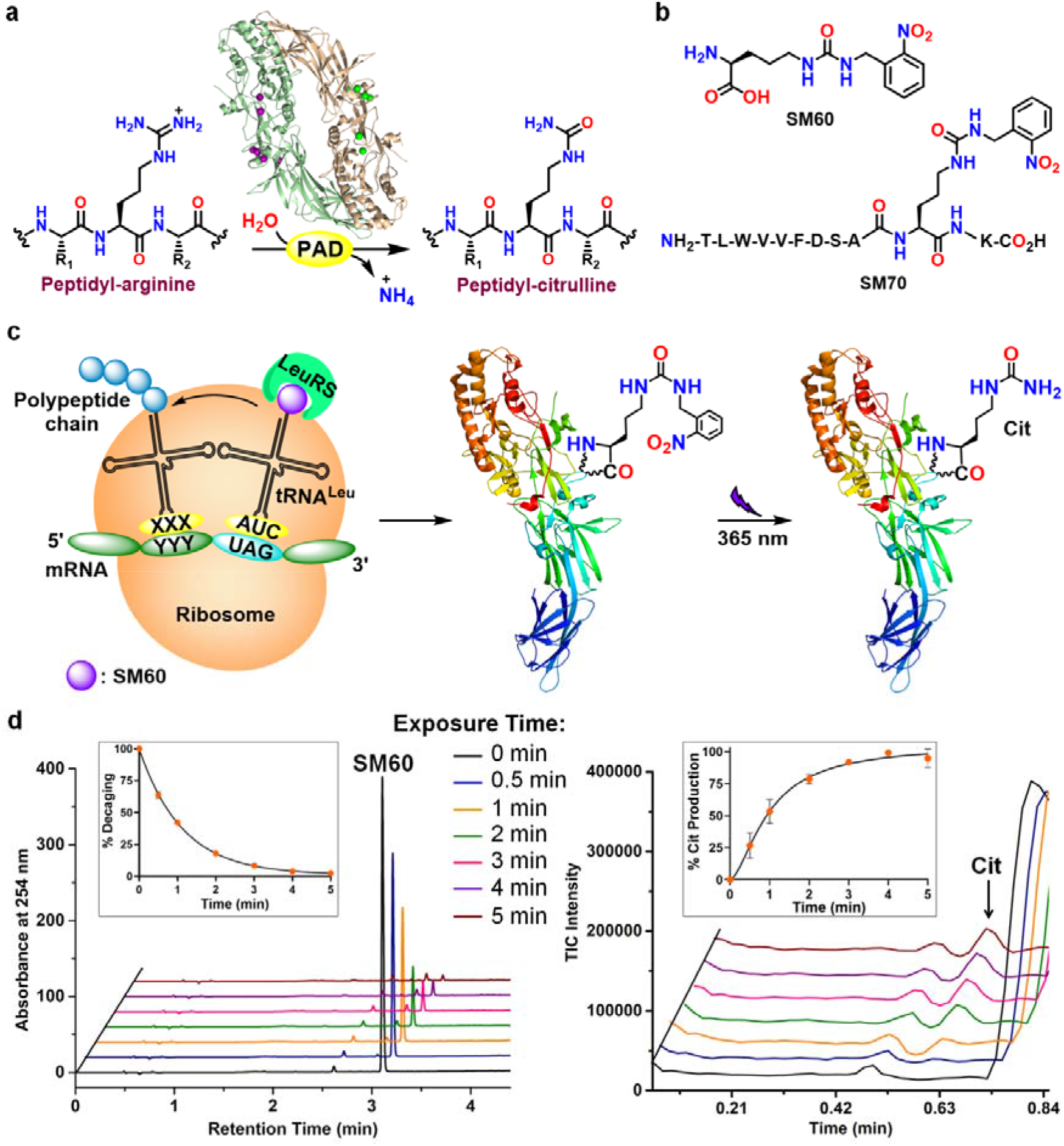
SM60, a photocaged-citrulline and its conversion to citrulline with 365 nm UV. **a**, Conversion of peptidyl-arginine to peptidyl-citrulline by the PADs. **b**, Chemical structures of **SM60** and **SM70**. **c**, Schematic representation of the incorporation of **SM60** into proteins by an engineered LeuRS-tRNA^Leu^ pair and subsequent conversion to citrulline. **d**, Decaging of **SM60** to citrulline with 365 nm UV irradiation. Left and right panels indicate the HPLC and ion chromatograms, showing the disappearance of **SM60** and the formation of citrulline (Cit), respectively, with increasing UV exposure. Quantitative analyses of decaging and Cit formation are shown in the insets. Assay mixture: 1 mM **SM60**, 2 mM DTT, Phosphate-buffered saline pH 7.4.

Aberrant protein citrullination is a hallmark of multiple autoimmune disorders, including rheumatoid arthritis (RA), multiple sclerosis (MS), ulcerative colitis (UC) and lupus, as well as several neurodegenerative diseases and cancer. Of note, multiple pan- and isozyme-selective PAD inhibitors are known and these inhibitors show efficacy in animal models of RA, UC, MS, lupus and sepsis.^1,8,10–12^ The contribution of protein hypercitrullination to the pathology of various diseases has been further established using the phenylglyoxal (PG)-based citrulline-specific probes, Rhodamine-PG (Rh-PG) and Biotin-PG.^1,8^ For example, Rh-PG enabled visualization of extensive citrullination of serum proteins and a marked decrease upon treatment with pan-PAD inhibitor, Cl-amidine, in a mouse model of UC.^13^ Using Biotin-PG and a chemoproteomic platform, we also identified various classes of novel citrullinated proteins, including serine protease inhibitors (SERPINs), serine proteases, transport proteins and complement system components along with known citrullinated proteins (e.g., vimentin, enolase, keratin and fibrin) in the serum, synovial fluid and synovial tissue of RA patients.^14^ Although the list of citrullinated proteins is ever expanding, the effect of citrullination on the structure and activity of a given protein remains poorly understood.

Currently, the most commonly used strategy for generating a citrullinated protein involves its treatment with a PAD. However, this leads to citrullination at all sites that are available *in vitro*, which may not fully recapitulate the situation *in vivo*. Moreover, the degree of modification at each site is frequently partial, leading to a complex heterogeneous mixture.^14–16^ Clearly, this strategy fails to provide information on the effect of individual citrullination events, underscoring the need for a method to site-specifically incorporate citrulline into proteins.

Although Gln mutations have been used as surrogates for citrulline (Cit),^17^ Gln is smaller and does not accurately mimic the H-bonding patterns afforded by Cit. *In vitro* translation systems or post-translational mutagenesis approaches that have been used to incorporate Cit are limited by their cumbersome nature, the need for specialized equipment, and for the latter approach, the need to incorporate a dehydroalanine at the site of modification, which is itself challenging and generates a mixture of D- and L-stereoisomers.^18,19^ Additionally, these strategies preclude the expression of site-specifically citrullinated proteins in living cells, and therefore, are ineffective for interpreting the downstream implications of this PTM. By contrast genetic code expansion technologies enable the site-specific incorporation of unnatural amino acids (UAAs) into proteins using engineered aminoacyl-tRNA synthetase (aaRS)-tRNA pairs.^20–24^ This technology has been used to genetically encode many important PTMs, enabling the expression of homogeneously modified protein at desired sites in living cells.^25–28^ However, genetically encoding Cit using this technology has remained elusive so far.

In this paper, we report the facile site-specific incorporation of Cit into proteins in mammalian cells using an *E. coli*-derived engineered leucyl-tRNA synthetase (EcLeuRS)-tRNA_CUA_^EcLeu^ pair. This pair, in response to a nonsense codon (UAG), charges a photocaged-citrulline, **SM60** (Figure 1b, c), into proteins expressed in HEK293T or EXPI293F cells. Subsequently, the photocage is removed with 365 nm UV to generate Cit *in vitro* or in living cells (Figure 1c). To demonstrate proof-of-concept, we incorporated citrulline (Cit) at two well-known autocitrullination sites, R372 and R374, in PAD4 and elucidated how these modifications impact enzyme activity.

## Results

### Development of SM60: a Photocaged-Citrulline

Envisioning that it may be challenging to develop an engineered aaRS that would selectively charge Cit, while discriminating against a nearly isostructural arginine, we hypothesized that these challenges could be overcome through a caging strategy. As such, we designed a photocaged-citrulline (**SM60**, comprising an o-nitrobenzyl photocage on the Cit side chain), which is structurally distinct from the canonical amino acids but can be efficiently converted to Cit post-translationally (Figures 1b, C). **SM60** was synthesized over two steps from L-ornithine, and was characterized by ^1^H, ^13^C NMR spectroscopy and mass spectrometry (Figures S1-2). Using LC-MS studies, we found that **SM60** can be quantitatively converted to Cit in phosphate-buffered saline (PBS) supplemented with dithiothreitol (DTT) using 365 nm UV radiation for 5 min (Figures 1d, S3). Quantitative conversion was further supported by ^1^H NMR analysis, which shows the rapid disappearance of the benzylic protons of **SM60** at 4.5 ppm with increasing UV exposure and by the photodecaging of **Fmoc-SM60** to **Fmoc-Cit** (Figures S4-5). To further investigate the feasibility of decaging **SM60** on proteins, we synthesized a **SM60**-containing peptide, **SM70** (Figures 1b, S2), that contains residues 363-372 of PAD4 with **SM60** at the 372 position, a known autocitrullination site (see below). Gratifyingly, **SM70** also undergoes photodecaging to form the citrulline-containing peptide (Figure S6), indicating that **SM60** can be decaged in the presence of other amino acids.

### Genetically encoding SM60 in eukaryotes

Four different aaRS/tRNA pairs have been successfully engineered for incorporating UAAs in eukaryotic cells: bacteria-derived tyrosyl, tryptophanyl, and leucyl pairs and the archaea-derived pyrrolysyl pair.^21–24,29,30^ The first two pairs are restricted to structural analogs of phenylalanine and tryptophan, respectively, precluding their use to genetically encode **SM60**. However, both the archaeal pyrrolysyl (PylRS/tRNA^Pyl^) and *E. coli* leucyl (EcLeuRS-tRNA_CUA_^EcLeu^) pairs have been engineered to charge UAAs structurally similar to **SM60**. Engineered aaRSs often exhibit substrate polyspecificity, i.e., the ability to use several structurally analogous UAAs, while discriminating against the canonical amino acids. This property has provided a facile route to rapidly expand the repertoire of genetically encoded UAAs without having to engineer new aaRS mutants for each distinct substrate. To explore if such a polyspecific aaRS can accept **SM60** as a substrate, we screened several existing PylRS and EcLeuRS mutants using an EGFP-39-TAG expression assay in HEK293T cells in the presence of their cognate amber suppressor tRNA. This screen identified an EcLeuRS mutant (M40I, Y499I, Y527A and H529G in the active site, and T252A in the editing domain)^31^ which enabled robust expression of the fluorescence reporter only when **SM60** was supplemented in the medium (Figure 2a,b). Purification of the resulting full-length EGFP using a C-terminal polyhistidine tag, followed by mass spectrometry, showed a mass consistent with the successful incorporation of **SM60** (Figures 2c,d). Furthermore, UV irradiation of cells expressing EGFP-39-**SM60** before lysis followed by protein purification and MS analysis afforded a single protein mass consistent with the complete deprotection and incorporation of citrulline at position 39 of EGFP (Figure 2d).

**Figure 2.**
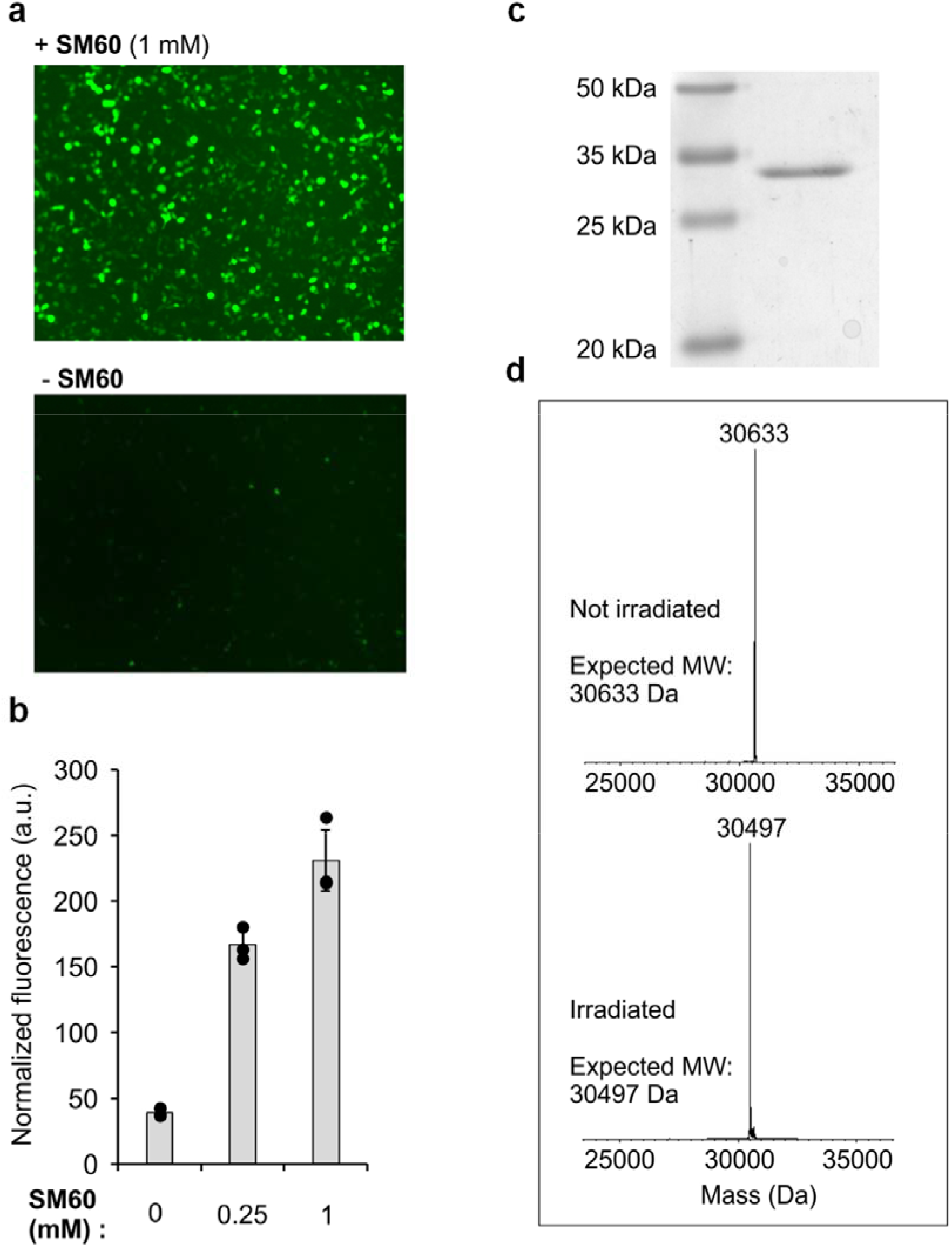
Site-specific incorporation of SM60 in EGFP and subsequent conversion to citrulline in HEK293T cells. **a**, EGFP-39-TAG reporter expression by EcLeuRS-tRNA_CUA_^EcLeu^ pair in HEK293T cells in the presence of **SM60** indicated by the fluorescence of EGFP. **b**, Quantification of EGFP-39-TAG reporter expression efficiency in the presence of increasing concentration of **SM60**. **c**, Coomassie stain of purified EGFP containing **SM60**. Full gel is given in Figure S7. **d**, Deconvoluted mass spectrum of EGFP before and after 365 nm UV irradiation, indicating the presence of **SM60** and citrulline, respectively, at 39 position. Non-deconvoluted spectra are given in Figures S8-9.

### Site-specific Incorporation of Citrulline in PAD4

Having established our ability to site-specifically incorporate Cit into EGFP, we sought to exploit this technology to address the effect of autocitrullination on PAD4 activity. We focused on these studies because the effect of autocitrullination on PAD4 activity has been debated. While Andrade *et al.* reported that autocitrullination negatively impacts PAD4 activity, we showed that autocitrullination has little to no impact on PAD4 activity.^17,32^ Using a citrulline-specific fluorescent probe Rh-PG,^13^ we confirmed that PAD4 autocitrullinates in the presence of Ca^+2^ in a time-dependent manner (Figure 3a). We and others have previously mapped several autocitrullination sites in PAD4 (Table S1 and Figure S10).^17,32^ While most of these sites are on the surface, the frequently observed R372 and R374 sites are present in the active site. Notably, the guanidinium groups on these two residues are only 3.5 Å from each other and the expected electrostatic repulsions are delicately balanced by H-bonding and salt-bridge interactions with D345. Moreover, R374 forms two H-bonds with the small molecule substrate, BAA.^33^ Therefore, we expected that citrullination at these sites would significantly impact enzyme activity. To evaluate this hypothesis, we incorporated Cit at positions 372 and 374 in PAD4. Wild-type (WT) PAD4 and the 372 and 374 TAG mutants were separately cloned into a pAcBac3 plasmid,^31,34^ which also encodes the mutant EcLeuRS and 8 copies of the tRNA ^EcLeu^. Subsequently, these plasmids were transfected separately into HEK293T cells, and the PAD4 protein or its mutants (after irradiation to remove the photocage), were purified using a C-terminal polyhistidine tag. While WT PAD4 expression was robust (10 μg/10^7^ cells), yields for the mutants were very low, indicating poor suppression efficiency at these sites. The Chin group has recently reported a mutant eukaryotic release factor (eRF1 E55D) that can enhance TAG-suppression efficiency in mammalian cells upon overexpression.^35^ To explore if this strategy can overcome the low suppression efficiency at the target sites in PAD4, we cloned eRF1-E55D mutant under a CMV promoter in a pIDTSMART vector. Indeed, co-transfection of this plasmid significantly improved the efficiency of nonsense suppression and enabled the purification of the desired mutants (2-4 μg/10^7^ cells, Figure 3b,c). Since the +0.98 Da mass change upon citrullination is difficult to detect by intact MS analysis, we confirmed the incorporation of citrulline at the desired position by LC-MS/MS analysis of the peptides resulting from Lys-C/Glu-C digestion of PAD4 (Figure S11 and Table S2). Using a similar procedure, we also expressed and purified wild-type PAD4 and the R374Cit mutant from EXPI293F cells that grow in suspension (Figure S12).

**Figure 3.**
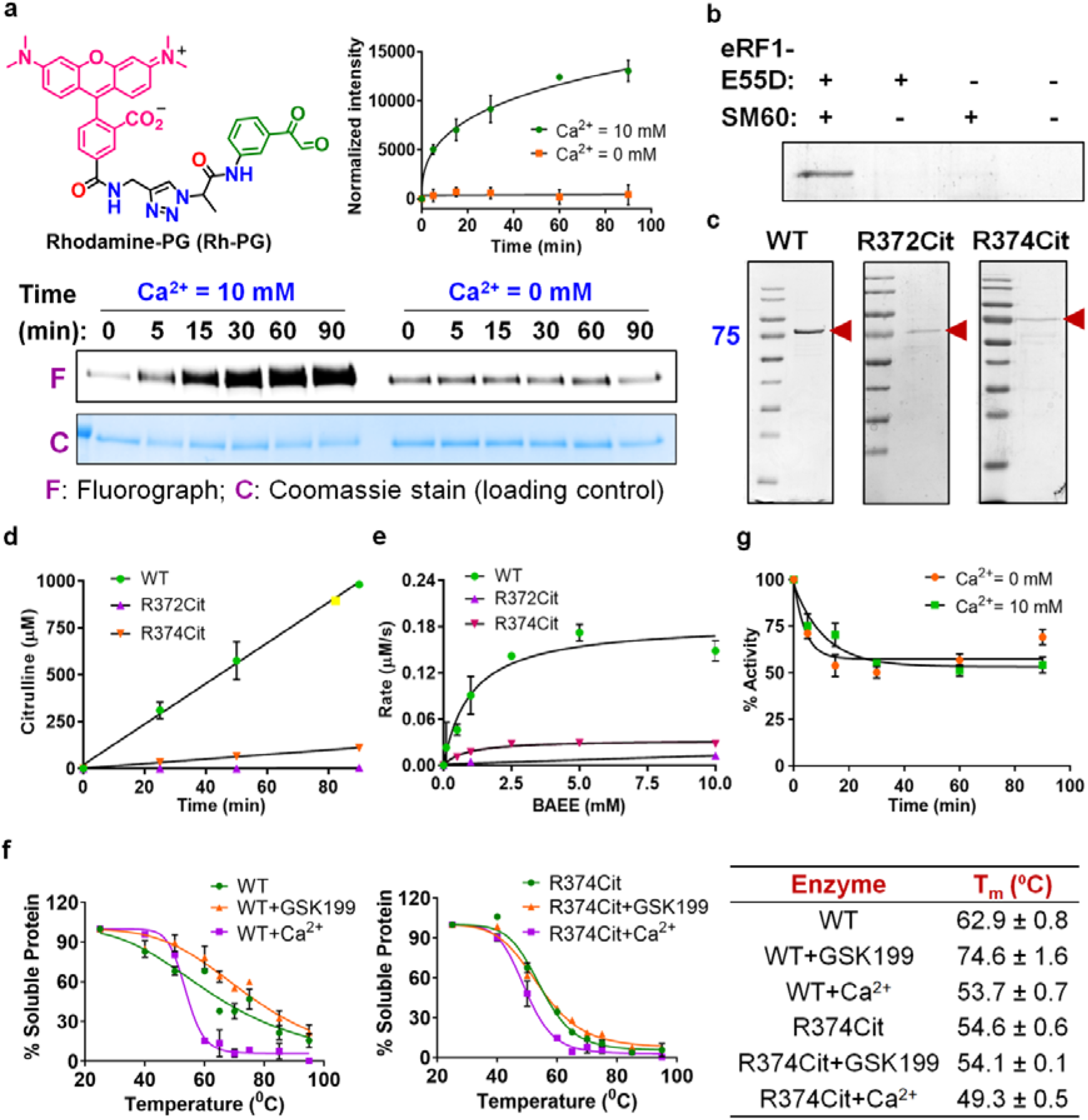
Autocitrullination of PAD4, incorporation of citrulline in PAD4 and implications thereof. **a**, Chemical structure of Rh-PG and the fluorescence labeling of autocitrullinated PAD4. The bands at 0 min in the presence of calcium and at all the time-points in the absence of calcium correspond to the basal levels of autocitrullination during the expression and purification of PAD4 from *E. coli*. **b**, Expression of R372Cit PAD4 in the presence of engineered release factor, eRF1-E55D and SM60 in HEK293T cells, indicating the essential role of eRF1-E55D for efficient TAG-suppression. **c**, Coomassie stains for WT, R372Cit and R374Cit PAD4, indicating their purity. **d**, Time-dependent citrulline production by WT, R372Cit and R374Cit PAD4. **e**, Michaelis-Menten kinetics for WT, R372Cit and R374Cit PAD4. **f**, Thermal shift profiles of WT and R374Cit PAD4. Western blot images are given in supplementary Figure S15. The table indicates the melting temperatures (T_m_). **g**, Effect of autocitrullination on the enzymatic activity of PAD4.

### Activity of WT, R372Cit, R374Cit and Autocitrullinated PAD4

We first ensured that the biochemical activity and calcium dependence of WT PAD4 expressed from HEK293T cells (PAD4_Mam_) and *E. coli* (PAD4_Bac_) are similar (Figure S13 and Table S3). Next, we determined kinetic values for the R372Cit and R374Cit mutants. Notably, both WT and the R374Cit mutant exhibited a time-dependent increase in citrulline production. By contrast, the R372Cit mutant produced only a negligible amount of citrulline after 90 min (Figure 3d). Furthermore, the rate of citrulline production, indicated by the slope of the straight line, is significantly lower for the R374Cit mutant than WT PAD4. In agreement, the *k*_cat_/*K*_m_ values of the R374Cit and R372Cit mutants are 9- and 181-fold lower than that for WT PAD4 (Figure 3e and Table S3).

Intrigued by these results, we sought to understand why the activity of these Cit-containing mutants is lower than WT PAD4. Kinetic studies indicated that R374Cit mutant possesses a similar *K*_m_, but a 10-fold lower *k*_cat_ than WT PAD4 (Table S3), suggesting a slow conversion of substrate to product. Furthermore, RFA, a PAD-targeted activity-based probe that covalently modifies the active site cysteine, C645,^36^ fluorescently labeled only WT PAD4 when tested with both purified enzymes and enzyme-containing cell lysates (Figure S14). Based on these observations, we hypothesized that citrullination may induce local conformational changes within the active site, leading to very slow or no reaction between C645 and the guanidium group of substrate, BAEE or the fluoroacetamidine warhead on RFA. To investigate this possibility, we performed a thermal shift assay in the presence of a PAD4-selective ligand, GSK199, which binds to an allosteric pocket near the active site and H-bonds to both D473 and H471.^37^ Using this assay, we found the melting temperatures (T_m_) of WT and R374Cit PAD4 to be 62.9 and 54.6 °C, respectively (Figures 3f, S15). Despite having a lower T_m_, the R374Cit mutant is as stable as WT PAD4 at 37 °C (Figure S16). As expected, GSK199 increased the T_m_ of WT PAD4, however, it did not increase the T_m_ of the R374Cit mutant (Figure 3f). These results indicate that GSK199 binds poorly to the R374Cit mutant, likely due to local conformational changes around the active site. However, the overall folding of the mutant is the same as WT since both the WT and R374Cit proteins exhibit a similar ∆T_m_ in the presence of calcium (Figure 3f).

### Quantitative Proteomics of PAD4 Autocitrullination

As discussed earlier, conflicting reports indicated that autocitrullination can either inactivate the enzyme or have no effect. Since our present results suggest that autocitrullination should decrease PAD4 activity, we regenerated autocitrullinated PAD4 by incubating the enzyme in the presence of 10 mM CaCl_2_. Consistent with our previous observations, autocitrullinated PAD4 exhibits similar activity to control PAD4 (incubated in the absence of CaCl_2_) (Figure 3g). The activity loss for both autocitrullinated and control PAD4 in this experiment is likely due to the oxidation of C645 over time. Given that autocitrullination does not decrease enzyme activity, but citrullination of R372 and R374 does, we asked two questions. Are R372 and R374 the preferred sites of autocitrullination? Also, what fraction of PAD4 gets autocitrullinated? If only a small fraction of PAD4 is autocitrullinated, then this process should not impact the activity of uncitrullinated enzyme.

To answer these questions, we took a quantitative proteomics approach. PAD4 was autocitrullinated for various times, and digested with Glu-C and Lys-C to maximize peptide coverage. The resulting peptides were then labeled with tandem mass tags (TMT) and were subjected to tandem mass analysis (Figure S17). Enzyme incubated in the absence of calcium was used as the negative control. From this analysis, we identified 13 unique citrullination sites on PAD4 (Figure 4a and Tables S1, 4). Notably, these exclude the previously reported R156, R205, R383, R609 and R639 and include two new sites, R650 and R651. Although both R372 and R374 residues show a time-dependent increase in citrullination, citrullination of arginines 212/218, 484/488/495 and 650/651 occurs at much higher rate. For example, the extent of citrullination at arginines 212/218 and 484/488/495 is significantly higher at 5 min than that at the 372 and 374 sites after 90 min (Figures 4a, b). These observations indicate that arginines 372 and 374 are not the preferred sites of autocitrullination. Additionally, none of the observed autocitrullination sites exhibited a marked decrease in the arginine-containing parent peptide levels, which indicates that only a minor fraction of PAD4 undergoes autocitrullination, further explaining its nominal impact on the enzyme activity. Nonetheless, these results showcase how this technology provides the ability to systematically characterize the behavior of individual citrullinated isoforms of any protein – both *in vitro* and in living cells – providing a powerful new approach to understand the biology of this PTM.

**Figure 4.**
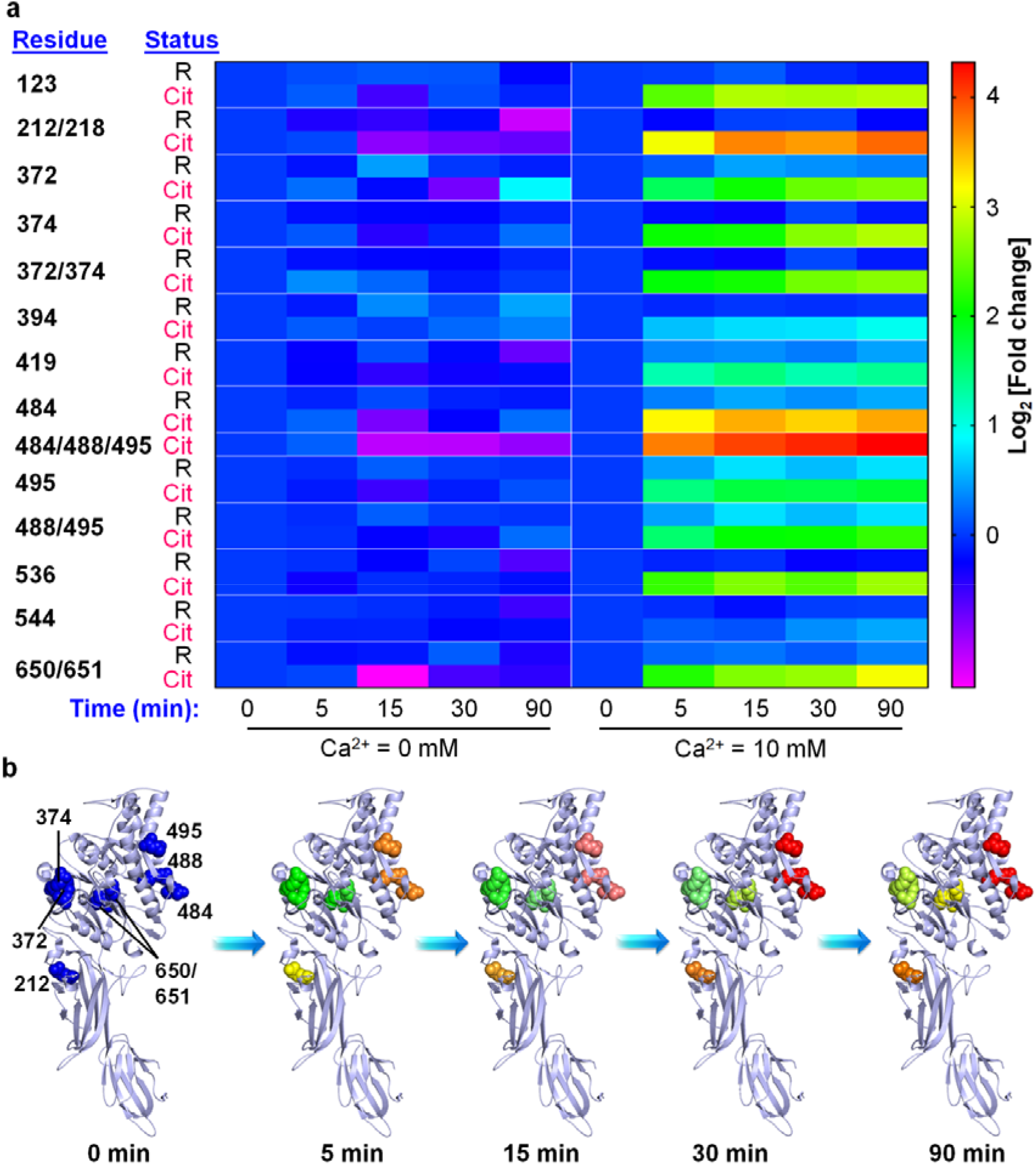
Autocitrullination sites in PAD4. **a**, Heat map representing the time-dependent change in peptides containing arginine or citrulline at the indicated positions (autocitrullination sites). Ca^2+^-untreated samples are negative controls. **b**, Time-dependent autocitrullination at the major sites. R218 site could not be shown because of disorder in that region (PDB: 1WDA).^33^ The increase in autocitrullination at these sites follows the same color code as in panel a.

## Discussion

Herein, we report the development of a novel technique for the site-specific incorporation of citrulline into proteins in mammalian cells. Central to this technology are a photocaged citrulline (**SM60**) and an *E. coli*-derived engineered leucyl-tRNA synthetase (EcLeuRS)/tRNA_CUA_^EcLeu^ pair that enables the incorporation of **SM60** into proteins in response to a TAG nonsense codon with high fidelity and efficiency. Subsequently, the photocage is removed with 365 nm light to generate citrulline. Using this technology, we incorporated citrulline at two known autocitrullination sites, R372 and R374, in PAD4. Kinetic studies indicate that the R374Cit and R372Cit mutants are 9- and 181-fold less active than WT PAD4. Detailed studies indicate that citrullination induces local conformational changes within the active site that leads to slow reaction between C645 and the guanidium group of substrate, the first step in the catalytic cycle. While these results indicate that citrullination of R372 and R374 would decrease PAD4 activity, we found that autocitrullination does not impact the enzymatic activity. Quantitative proteomics studies indicate that 212/218 and 484/488/495, and not 372 and 374, are the preferred sites of citrullination. While faster autocitrullination of arginines 212/218 and 484/488/495 is likely due to their residence at the surface of PAD4, upon citrullination, they may expose deeply buried autocitrullination sites by conformational changes. Efforts are currently under way to elucidate the effect of citrullination at these major sites, particularly the 484/488/495 residues because they are present at the interface of the head-to-tail PAD4 dimer that is known to alter enzymatic activity.

Since it is well established that citrullination is critical for many physiological processes, as well as in disease pathology, this new method will provide a direct and accessible approach to understand the biology of this PTM at the molecular level. For example, histone H3 citrullination at R26 leads to the transcriptional activation of more than 200 genes in estrogen receptor-positive breast cancer cells and inhibits the methylation of the neighboring K27 residue by 30,000-fold.^15^ However, the mechanism of negative crosstalk between these two PTMs remains poorly understood. Additionally, we recently showed that serine protease inhibitors (SERPINs), nicotinamide N-methyl transferase (NNMT), and pyruvate kinase isoform M2 (PKM2) are citrullinated in patients suffering from rheumatoid arthritis. Notably, the citrullination of the SERPINs and NNMT dramatically abolishes the enzymatic activity, while citrullinated PKM2 exhibits 2-3-fold higher activity than the WT enzyme.^14^ However, the underlying reasons behind such biochemical phenomenon are unclear. Finally, citrullination has been reported to impact neutrophil extracellular trap formation, pluripotency, and efficient elongation by RNA PolII but, again, the underlying mechanisms remain unclear.^1,8^ With our new enabling technology, it is now possible to incorporate citrulline on demand and mechanistically address how this PTM impacts these fundamental biological processes and pathways.

## Acknowledgement

This work was supported by in part by NIH grant R35 GM136437 (A.C.), institutional funds provided by the University of Massachusetts Medical School, and NIH grant R35 GM118112 (P.R.T.).

## Author contributions

S.M. designed and synthesized **SM60** and **SM70**, purified PAD4 from EXPI293F cells, characterized PAD4 mutants by MS/MS, performed enzyme kinetics, thermal shift assays, proteomics studies and analyzed the data, and wrote the manuscript. S.W. cloned the plasmids, optimized the expression system for citrulline incorporation in mammalian cells, expressed and purified EGFP, PAD4 and their mutants from HEK293T cells. Y.Z. screened for **SM60** incorporation activity. S.S. performed some of the cloning, mutation and cell culture studies. P.R.T. and A.C. conceived the idea of site-specific citrulline incorporation into proteins, designed the experiments and revised the manuscript.

## Data Availability

All the raw data are available upon request to the corresponding authors by email.

## Biological Materials

All the plasmids are available upon request to the corresponding authors by email.

## Supporting information

**Table S1.**
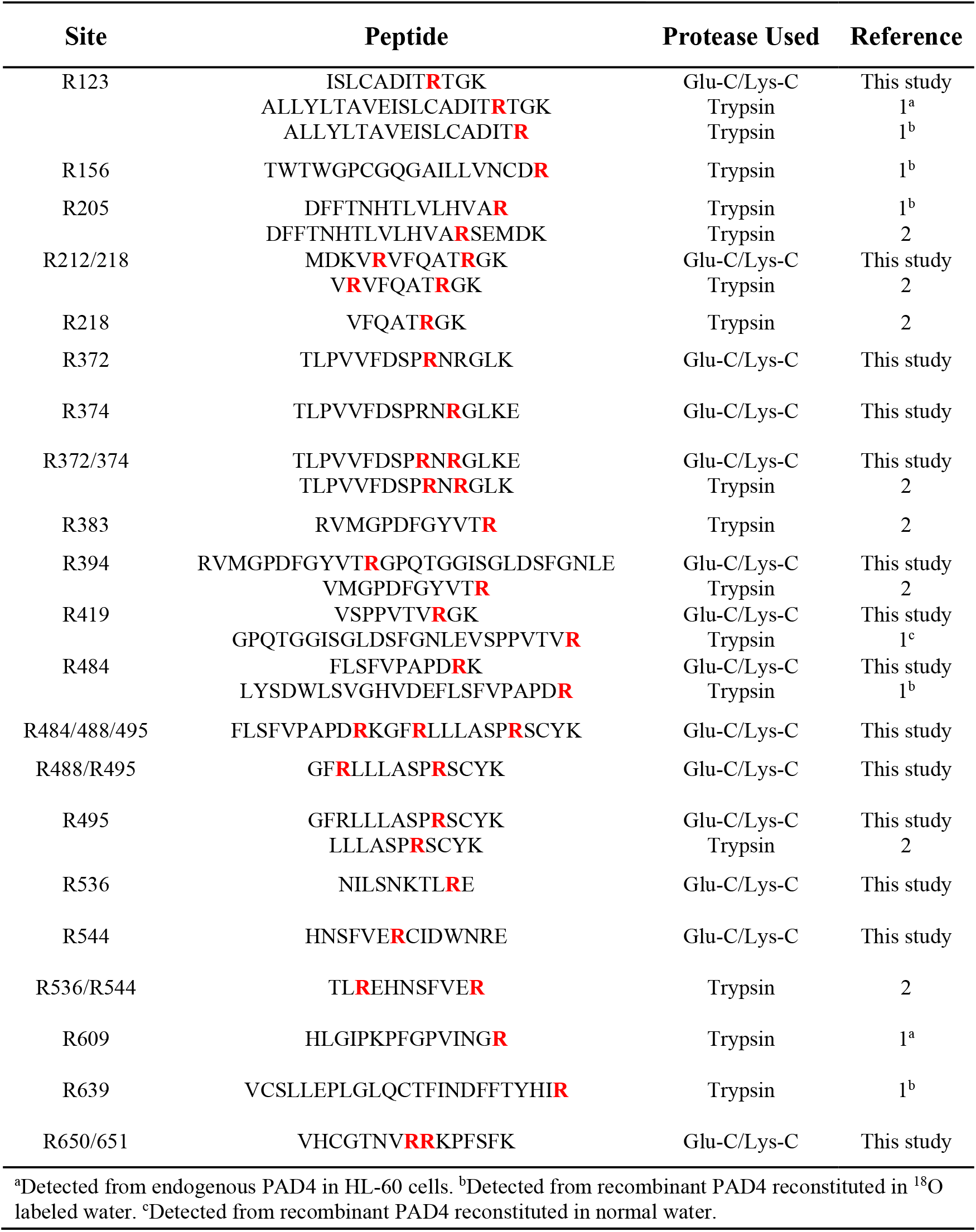
Sites of autocitrullination in PAD4 and peptides from which they are detected by tandem mass spectrometry.

**Table S2.**
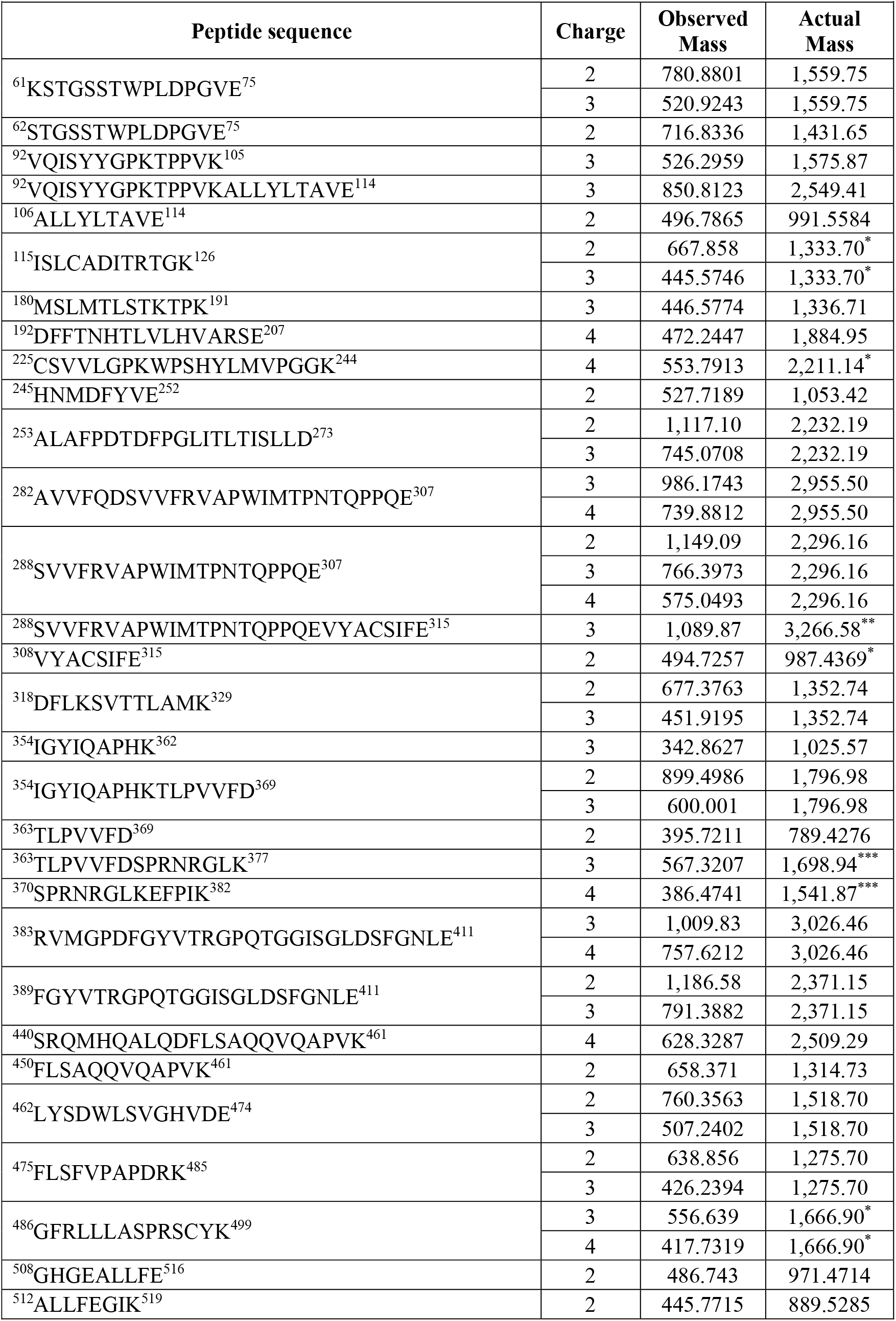
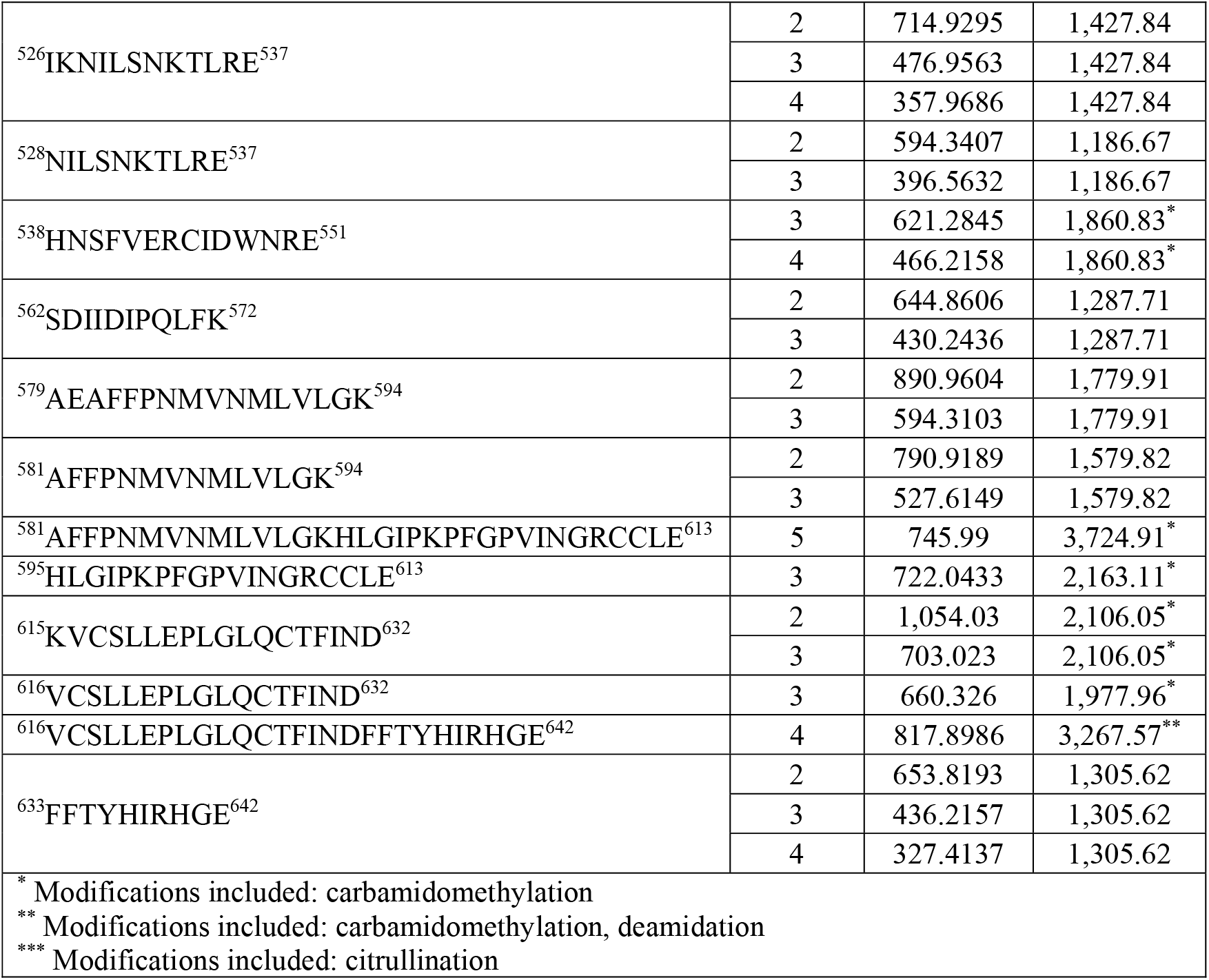
Ions Detected from LC-MS/MS Analysis of R374Cit PAD4 digested with Lys-C and Glu-C.

**Table S3.** Steady-state kinetic parameters for wild-type PAD4_Bac_, wild-type PAD4_Mam_, R372Cit and R374Cit mutants.

| Enzyme | $k_{\text{cat}}$<br>(S<sup>-1</sup>) | $K_{\text{m}}$<br>(mM) | $k_{\text{cat}}/K_{\text{m}}$<br>(M<sup>-1</sup>S<sup>-1</sup>) | $K_{0.5}$<br>(mM) |
| --- | --- | --- | --- | --- |
| PAD4<sub>Bac</sub> | 3.4 ± 0.2 | 0.7 ± 0.1 | 4740 ± 260 | 0.9 ± 0.10 |
| PAD4<sub>Mam</sub> | 3.7 ± 0.1 | 1.1 ± 0.2 | 3620 ± 810 | 0.9 ± 0.05 |
| R372Cit | - | - | 20 ± 2 | - |
| R374Cit | 0.33 ± 0.01 | 0.84 ± 0.06 | 390 ± 40 | - |

$k_{\text{cat}}$ : turnover number; $K_{\text{m}}$ : Michaelis-Menten constant; $k_{\text{cat}}/K_{\text{m}}$ : catalytic efficiency; $K_{0.5}$ : Ca<sup>2+</sup> concentration for half-maximal activity.

**Table S4.** Normalized^†^ fold changes of the peptides containing Arg and Cit at various autocitrullination sites with increasing time in the absence and presence of calcium.

|  |  |  |  |  |  |  |  |  |  |  |  |
| --- | --- | --- | --- | --- | --- | --- | --- | --- | --- | --- | --- |
| Site | Status | CaCl<sub>2</sub> (0 mM) |  |  |  |  | CaCl<sub>2</sub> (10 mM) |  |  |  |  |
|  |  | 0 min | 5 min | 15 min | 30 min | 90 min | 0 min | 5 min | 15 min | 30 min | 90 min |
| 123 | Arg | 1 | 1.060198 | 1.099362 | 1.086233 | 0.847332 | 1 | 1.015601 | 1.108801 | 0.955283 | 0.90346 |
|  | Cit | 1 | 1.111622 | 0.681444 | 1.061178 | 0.931525 | 1 | 5.35171 | 7.061624 | 6.900336 | 7.226682 |
| 212/218 | Arg | 1 | 0.763923 | 0.717972 | 0.867539 | 0.447513 | 1 | 0.844928 | 1.023137 | 1.030968 | 0.845279 |
|  | Cit | 1 | 1.043454 | 0.542113 | 0.587774 | 0.614152 | 1 | 8.693879 | 13.17746 | 12.295 | 14.45336 |
| 372 | Arg | 1 | 0.883519 | 1.368884 | 0.993322 | 0.918658 | 1 | 1.112136 | 1.391525 | 1.316463 | 1.254982 |
|  | Cit | 1 | 1.176907 | 0.858565 | 0.584389 | 1.82134 | 1 | 3.093736 | 4.228072 | 5.637283 | 6.04189 |
| 374 | Arg | 1 | 0.875998 | 0.849096 | 0.846745 | 0.939523 | 1 | 0.869947 | 0.809442 | 1.046085 | 0.90485 |
|  | Cit | 1 | 1.095559 | 0.741919 | 0.930278 | 1.176091 | 1 | 4.169863 | 4.267329 | 6.175974 | 7.094331 |
| 372/374 | Arg | 1 | 0.875998 | 0.849096 | 0.846745 | 0.939523 | 1 | 0.869947 | 0.809442 | 1.046085 | 0.90485 |
|  | Cit | 1 | 1.294953 | 1.149787 | 0.90417 | 1.006956 | 1 | 4.018527 | 4.30695 | 5.81589 | 6.233317 |
| 394 | Arg | 1 | 0.901459 | 1.284909 | 1.065847 | 1.397647 | 1 | 0.942636 | 0.987145 | 0.965936 | 0.99424 |
|  | Cit | 1 | 1.107009 | 1.01607 | 1.156688 | 1.256723 | 1 | 1.528377 | 1.672493 | 1.699763 | 1.930088 |
| 419 | Arg | 1 | 0.827406 | 1.071773 | 0.860154 | 0.609628 | 1 | 1.271619 | 1.311302 | 1.23086 | 1.387031 |
|  | Cit | 1 | 0.832199 | 0.72951 | 0.803851 | 0.869545 | 1 | 2.361985 | 2.763826 | 2.367449 | 2.578741 |
| 484 | Arg | 1 | 0.92959 | 1.047052 | 0.926588 | 0.899794 | 1 | 1.242001 | 1.414214 | 1.310393 | 1.420436 |
|  | Cit | 1 | 1.138131 | 0.573024 | 0.833161 | 1.173919 | 1 | 9.210844 | 11.60494 | 10.33882 | 11.87619 |
| 484/488/495 | Cit | 1 | 1.123889 | 0.467056 | 0.470848 | 0.52912 | 1 | 13.48546 | 16.22335 | 17.79418 | 19.97329 |
| 495 | Arg | 1 | 0.95418 | 1.118062 | 1.007654 | 0.982457 | 1 | 1.381913 | 1.689582 | 1.505247 | 1.691262 |
|  | Cit | 1 | 0.903753 | 0.697533 | 0.91088 | 1.074749 | 1 | 2.757447 | 3.363586 | 3.294364 | 3.547166 |
| 488/495 | Arg | 1 | 0.95418 | 1.118062 | 1.007654 | 0.982457 | 1 | 1.381913 | 1.689582 | 1.505247 | 1.691262 |
|  | Cit | 1 | 0.972655 | 0.82932 | 0.768438 | 1.170128 | 1 | 2.909961 | 4.05865 | 4.153517 | 4.771138 |
| 536 | Arg | 1 | 0.965267 | 0.835088 | 1.035026 | 0.652176 | 1 | 0.932817 | 0.970859 | 0.828171 | 0.870752 |
|  | Cit | 1 | 0.808321 | 0.960595 | 0.93045 | 0.872363 | 1 | 4.808432 | 6.058245 | 5.274009 | 6.644835 |
| 544 | Arg | 1 | 0.997692 | 0.963707 | 0.905216 | 0.692555 | 1 | 0.960062 | 0.872081 | 1.013397 | 1.022755 |
|  | Cit | 1 | 0.924236 | 0.929161 | 0.844401 | 0.875796 | 1 | 1.113087 | 1.076738 | 1.301342 | 1.410298 |
| 650/651 | Arg | 1 | 0.872161 | 0.893992 | 1.115739 | 0.769148 | 1 | 1.148698 | 1.202191 | 1.109467 | 1.248331 |
|  | Cit | 1 | 1.038139 | 0.379893 | 0.671116 | 0.728681 | 1 | 4.505436 | 6.083915 | 6.520579 | 8.805063 |
| <sup>†</sup>Ratios for the calcium- treated and untreated data set were normalized against the respective 0 min time-point. |  |  |  |  |  |  |  |  |  |  |  |

**Figure S1.**
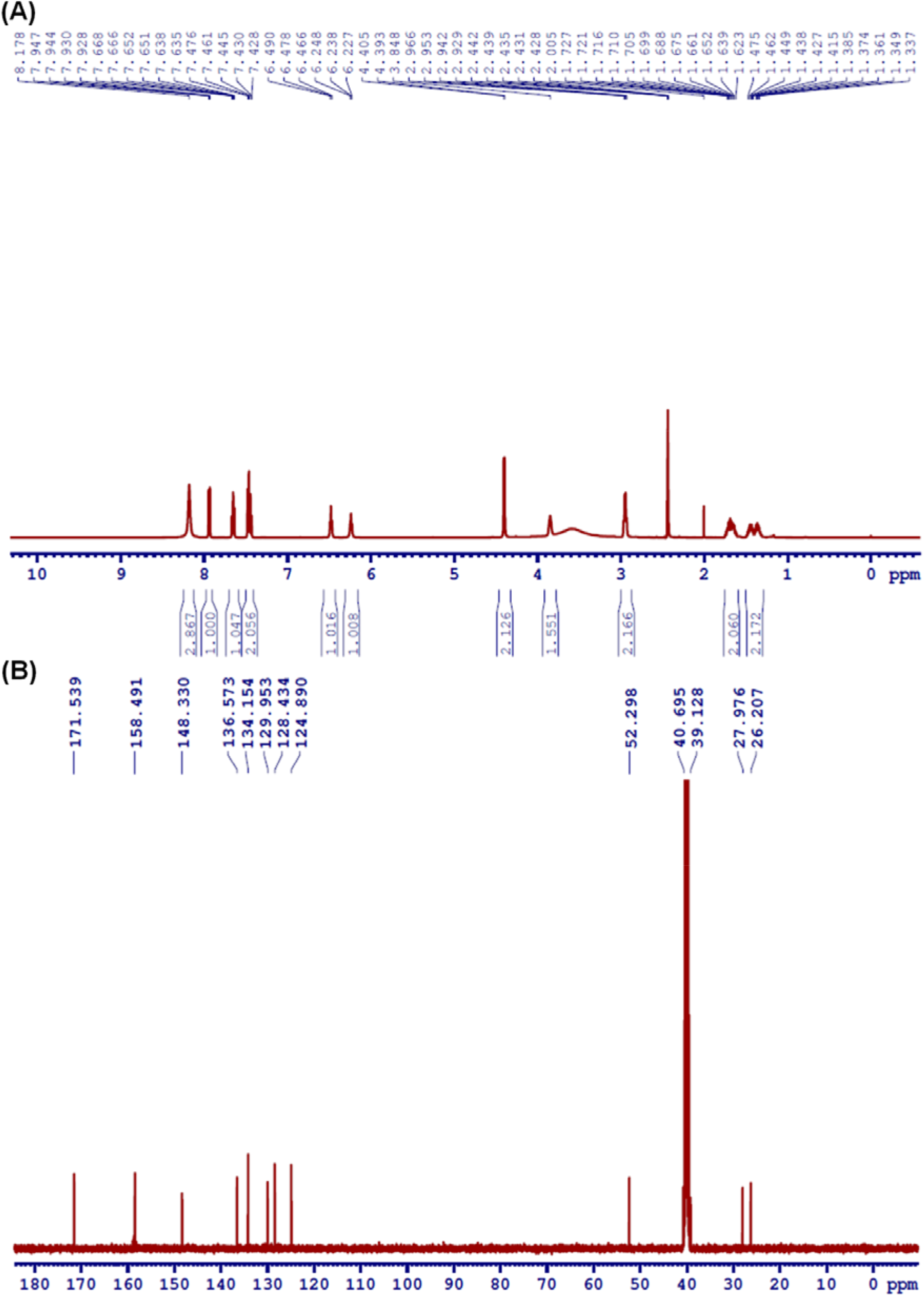
^1^H (A) and ^13^C (B) NMR of **SM60** in DMSO-*d_6_*.

**Figure S2.**
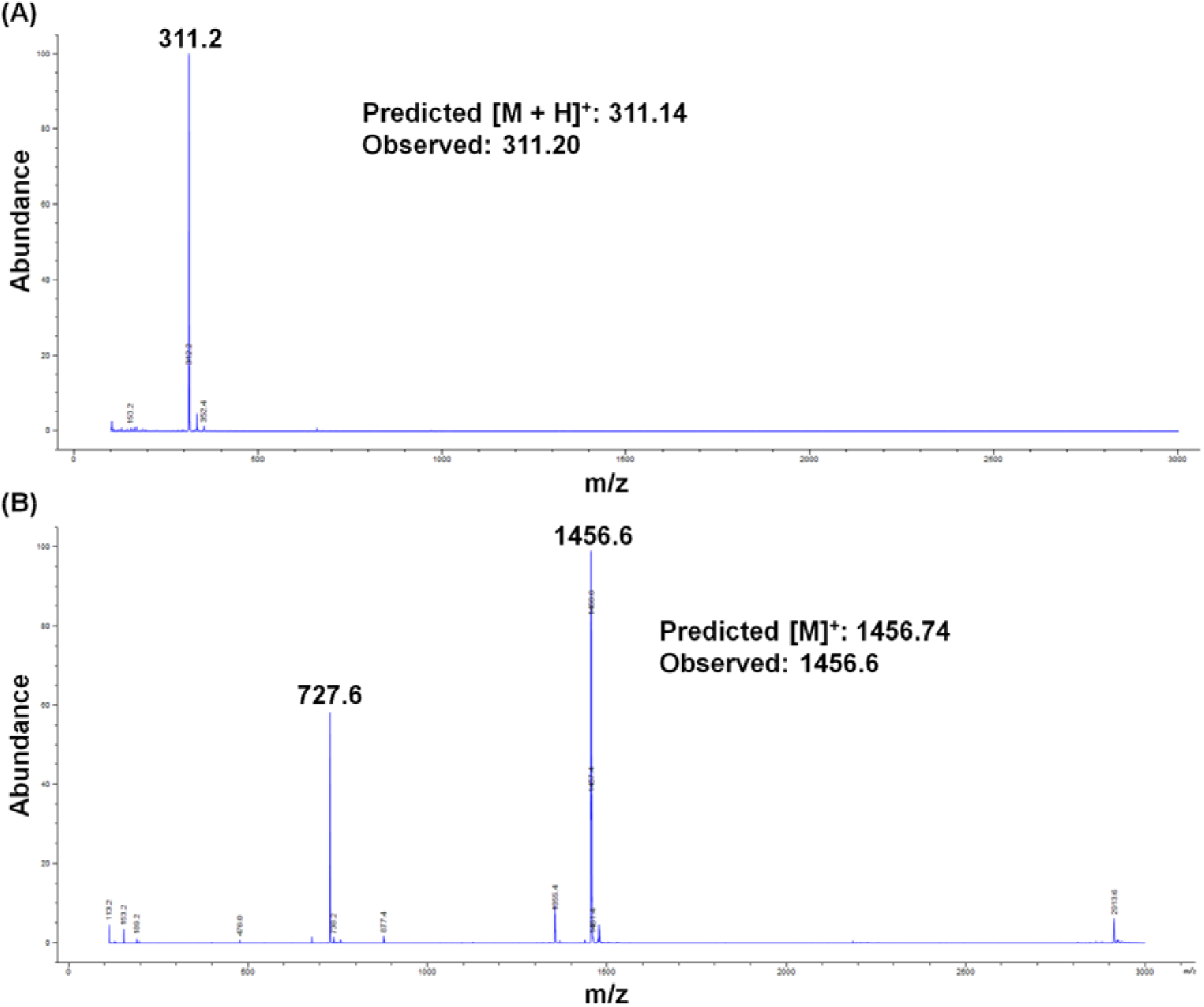
ESI Mass spectrum of **SM60** (A) and **SM70** (B).

**Figure S3.**
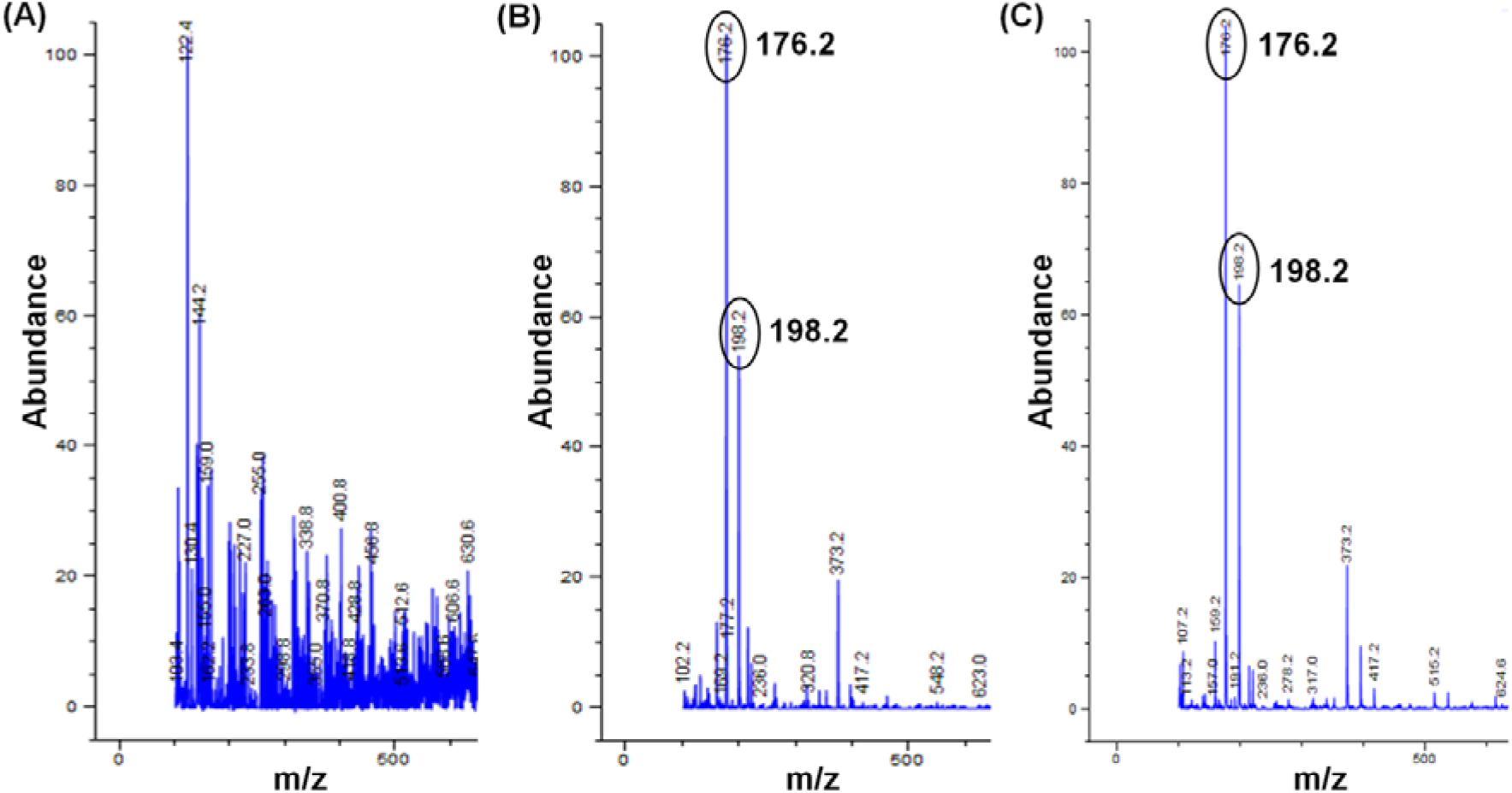
ESI mass spectrum for the formation of citrulline from **SM60** upon UV exposure. Citrulline appears at ~0.6 min in the ion chromatogram (Figure 1D). While the peaks for citrulline are absent before UV exposure (A), they appear upon UV exposure and reach a maximum after 5 min treatment. The representative mass spectrum after 5 min exposure is shown in panel B. (C) ESI mass spectrum of a standard sample of citrulline. ESI-MS (m/z) calculated for C_6_H_13_N_3_O_3_ [M + H]^+^: 176.10, found 176.20; [M + Na]^+^: 198.09, found 198.20.

**Figure S4.**
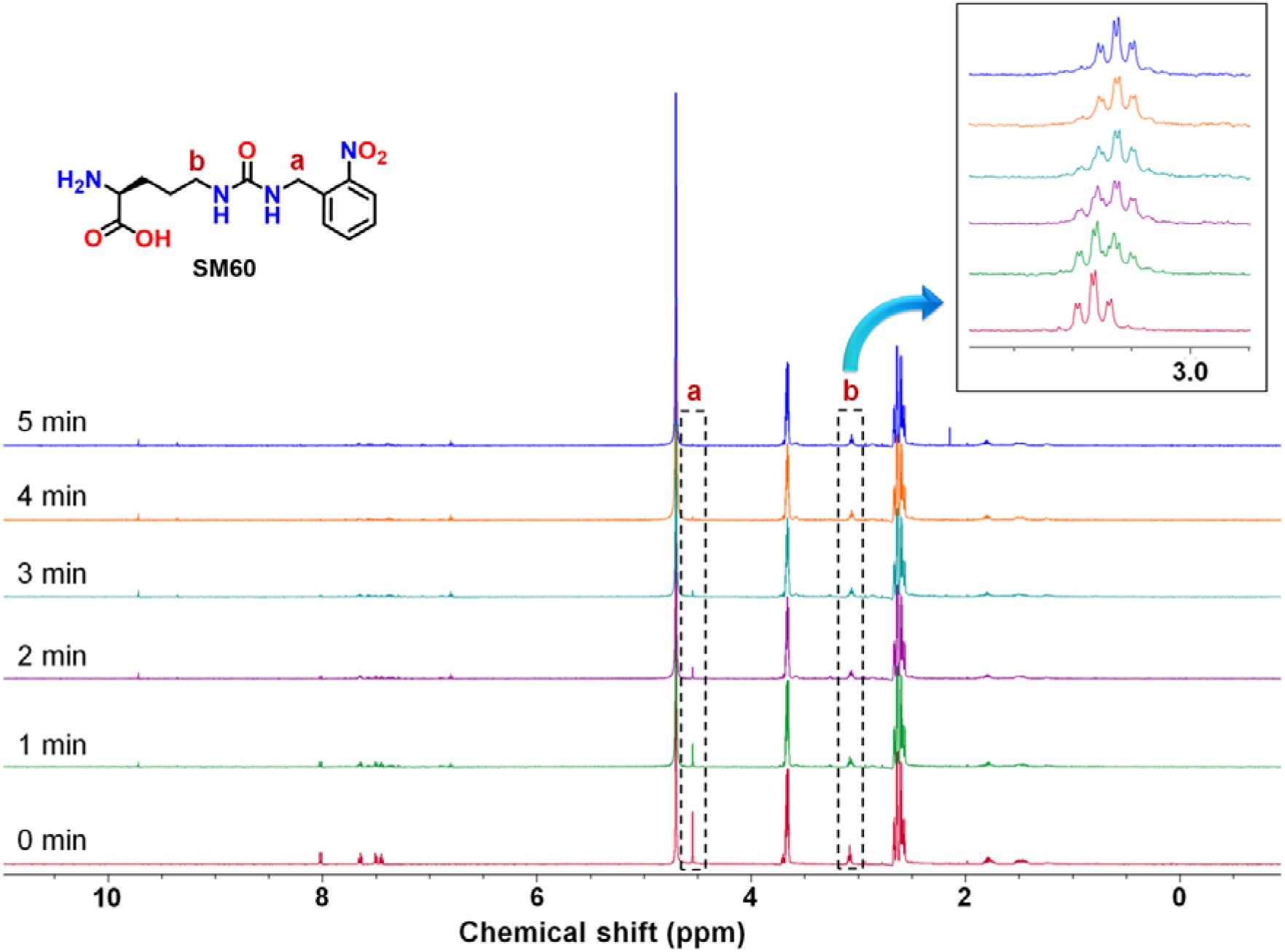
^1^H NMR spectrum of **SM60** in D_2_O after UV (365 nm) exposure for 1, 2, 3, 4 and 5 min. The disappearance of benzylic ‘a’ protons at 4.5 ppm (as seen in the spectrum denoted by 0 min) with increasing UV exposure indicates the complete removal of *o*-nitrobenzyl photocage. Furthermore, an upfield shift was also observed for the ‘b’ protons upon the formation of citrulline. The multiplets at 3.6-3.7 and 2.6-2.7 ppm are due to DTT used in the assay. Assay condition: 1 mM **SM60**, 2 mM DTT, D_2_O.

**Figure S5.**
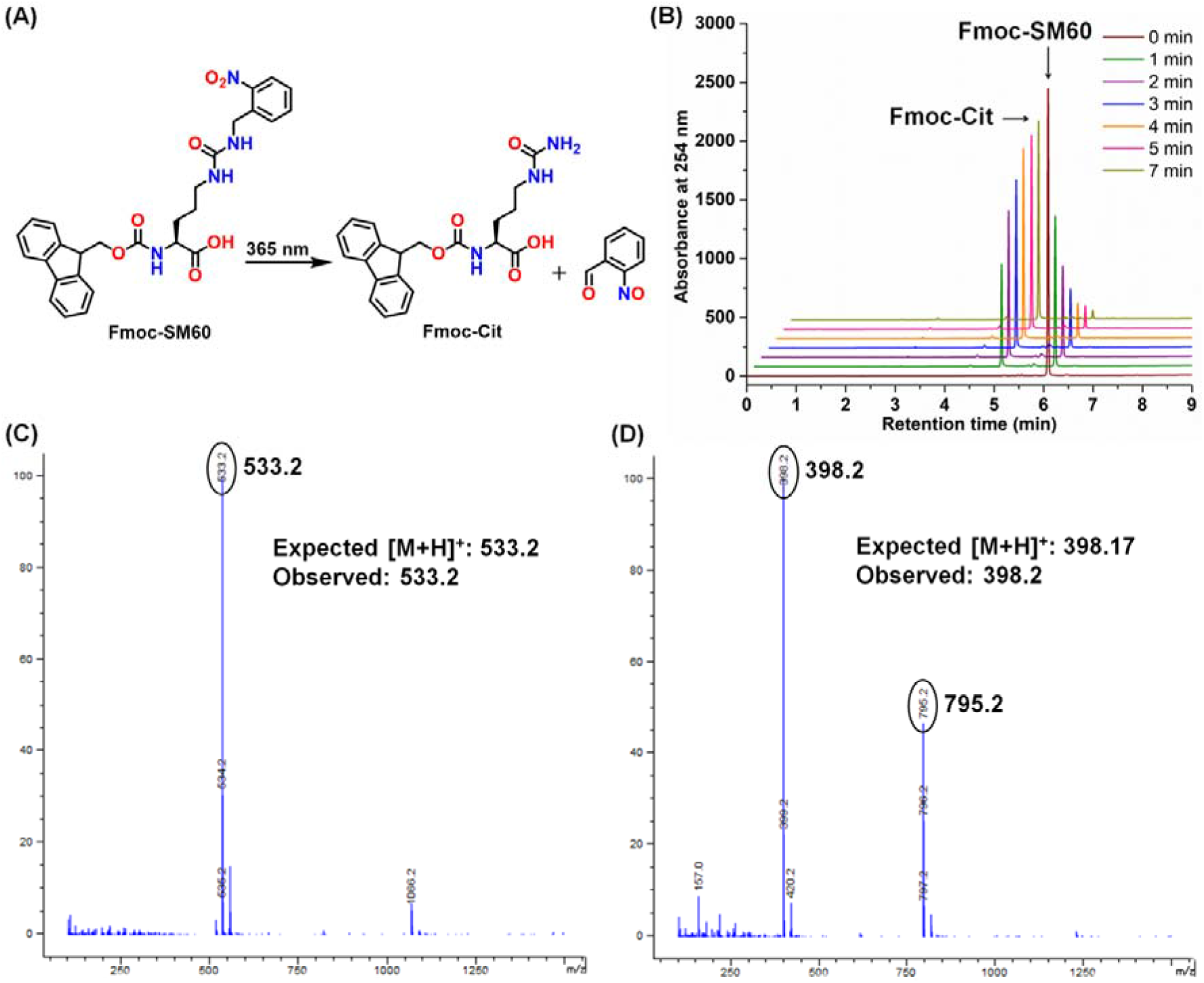
(A) Photodecaging of **Fmoc-SM60** to produce **Fmoc-Cit**. (B) HPLC chromatograms indicating the disappearance of **Fmoc-SM60** and the production of **Fmoc-Cit** with increasing UV exposure. These chromatograms also indicate the quantitative conversion of **Fmoc-SM60** to **Fmoc-Cit**. Assay mixture: 0.5 mM **Fmoc-SM60**, 2 mM DTT, Phosphate-buffered saline pH 7.4. ESI-Mass spectra of **Fmoc-SM60** (C) and **Fmoc-Cit** (D).

**Figure S6.**
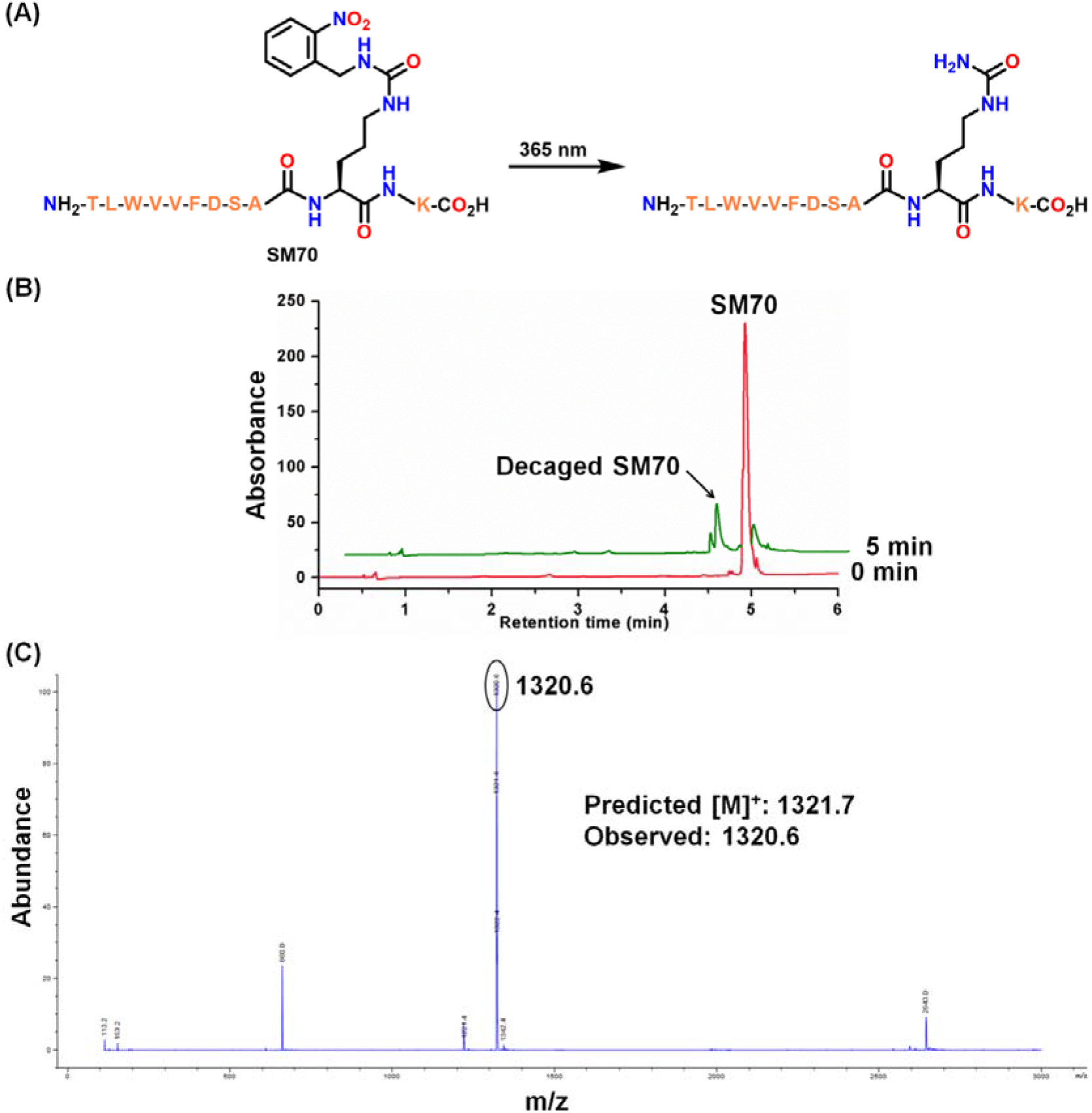
(A) Decaging of **SM70**, a PAD4-derived peptide (residues 363-372 with **SM60** at the 372 position. Prolines at 365 and 371 positions were replaced with W and A, respectively, and a lysine residue was added to the C-terminus for ease of synthesis.). (B) HPLC chromatogram of **SM70** before and after 5 min of 365 nm irradiation. The low peak intensity of the decaged product is explained by the lack of an aromatic group and its extremely low solubility in aqueous buffer. Even after performing the reaction in 1:1 acetonitrile/PBS, the solution became turbid upon the formation of decaged peptide. Assay condition: 200 μM **SM70**, 2 mM DTT, 1:1 acetonitrile/PBS (pH – 7.4). (C) ESI mass spectrum of the citrulline-containing peptide (decaged product).

**Figure S7.**
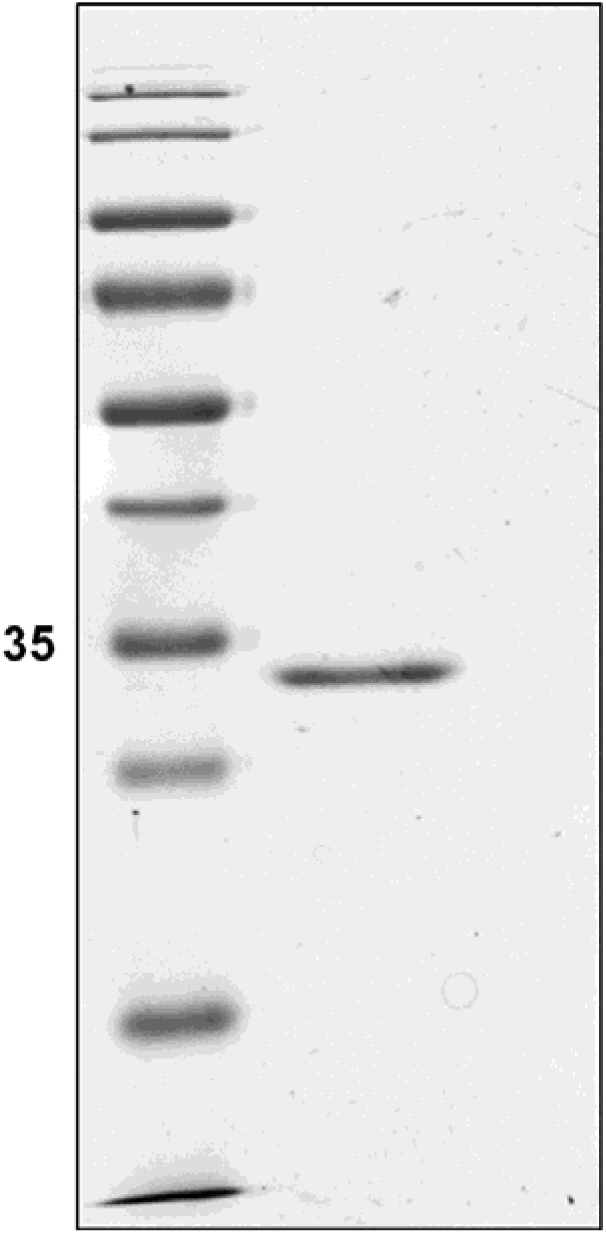
Coomassie stain of purified EGFP containing **SM60** at 39 position.

**Figure S8.**
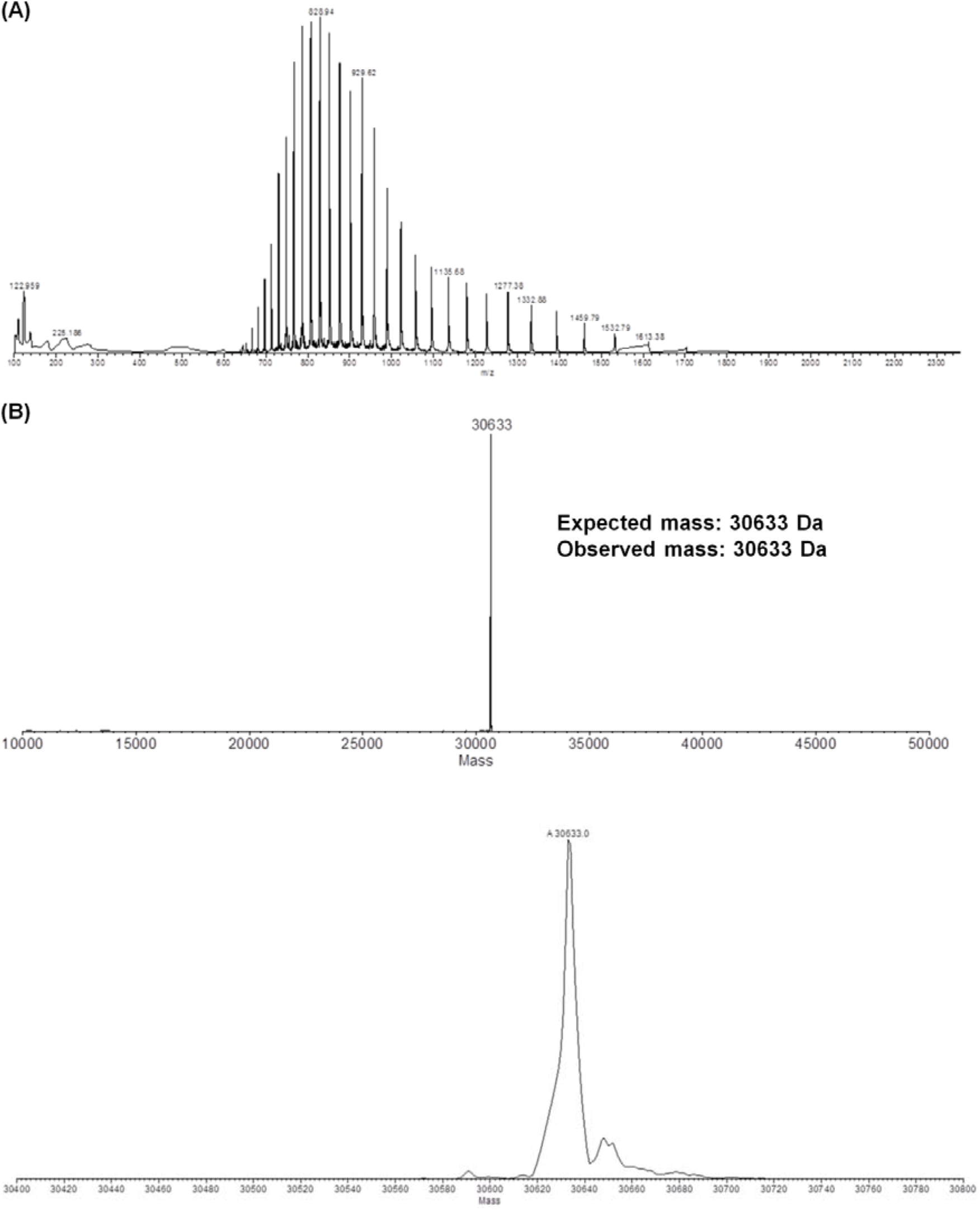
ESI mass spectrum (A) and deconvoluted spectra (B) of EGFP containing **SM60** at 39 position.

**Figure S9.**
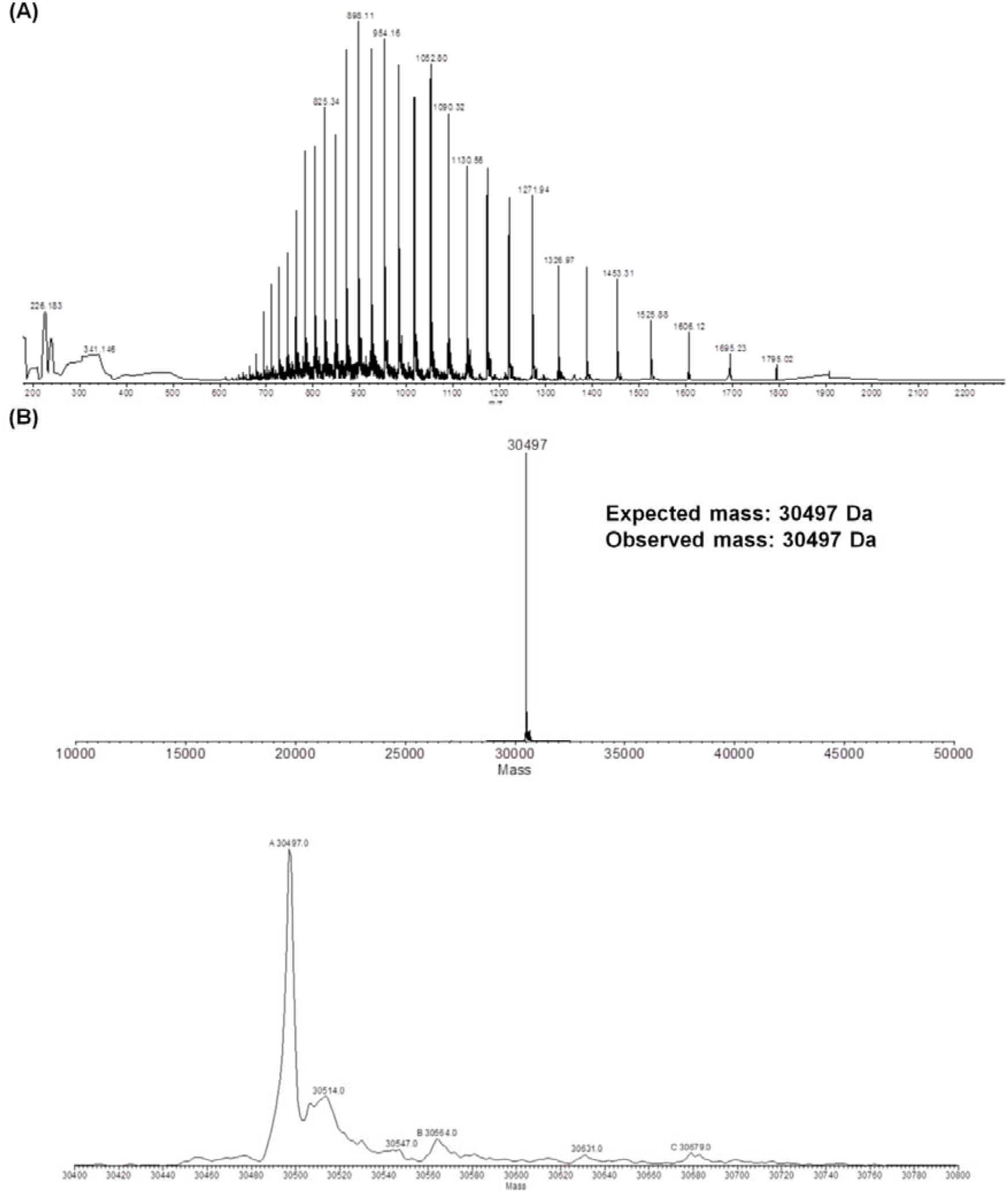
ESI mass spectrum (A) and deconvoluted spectra (B) of EGFP containing **Cit** at 39 position.

**Figure S10.**
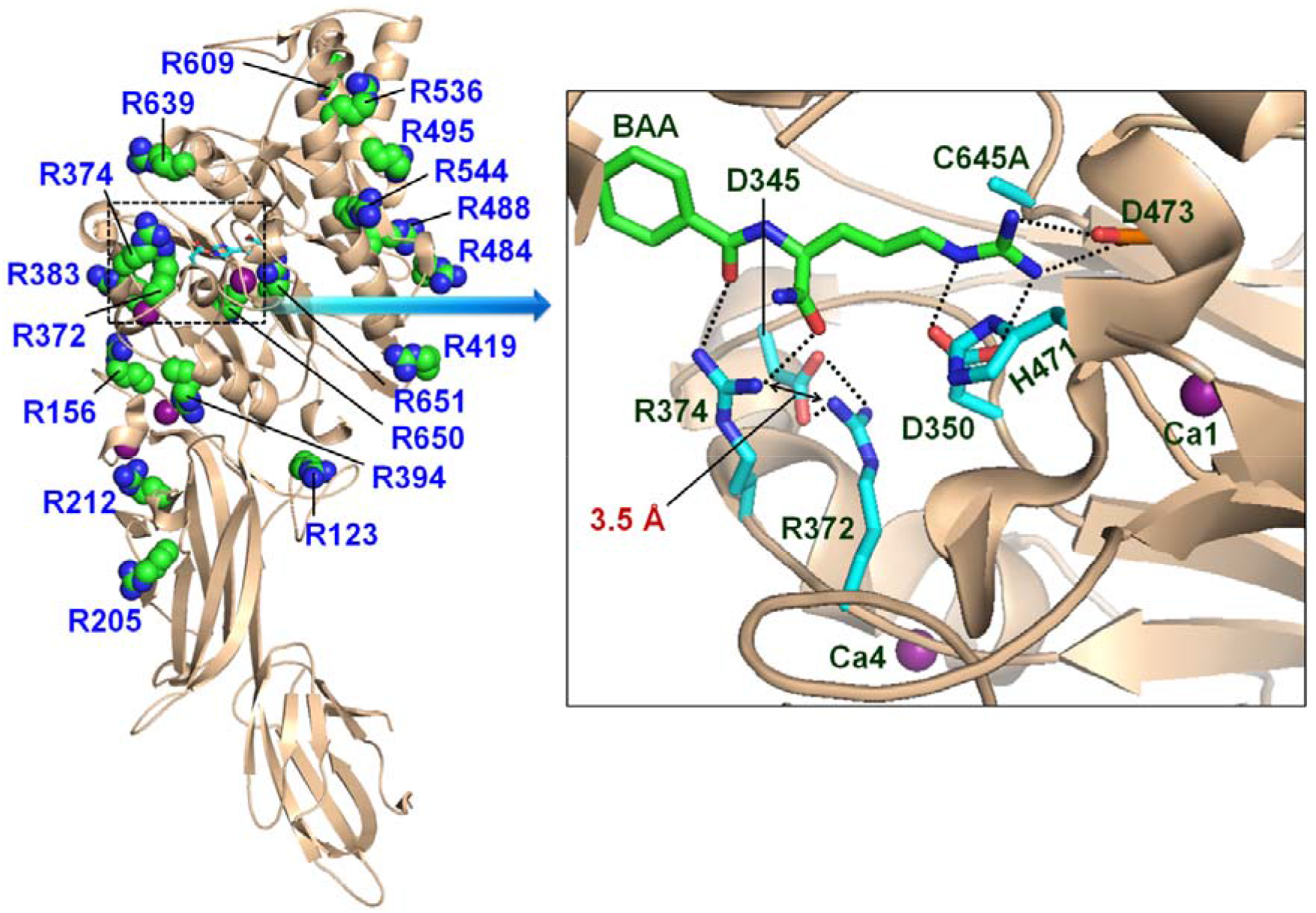
Sites of PAD4 autocitrullination (Table S1) (PDB code: 1WDA). R218 site could not be shown because of disorder in that region. While most of these sites are far from the active site and are on the surface of the protein, arginines 372 and 374 are close to the active site. Notably, the distance between these two positively-charged residues is only 3.5 Å, a value that is close to the sum of N∙∙∙N van der Waal’s radii (3.1 Å). Electrostatic repulsions between R372 and R374 are delicately balanced by hydrogen bonding interactions with the substrate, BAA and an aspartate, D345. These observations suggest that the citrullination of either of these residues will likely perturb the delicate balance and affect enzymatic activity.

**Figure S11.**
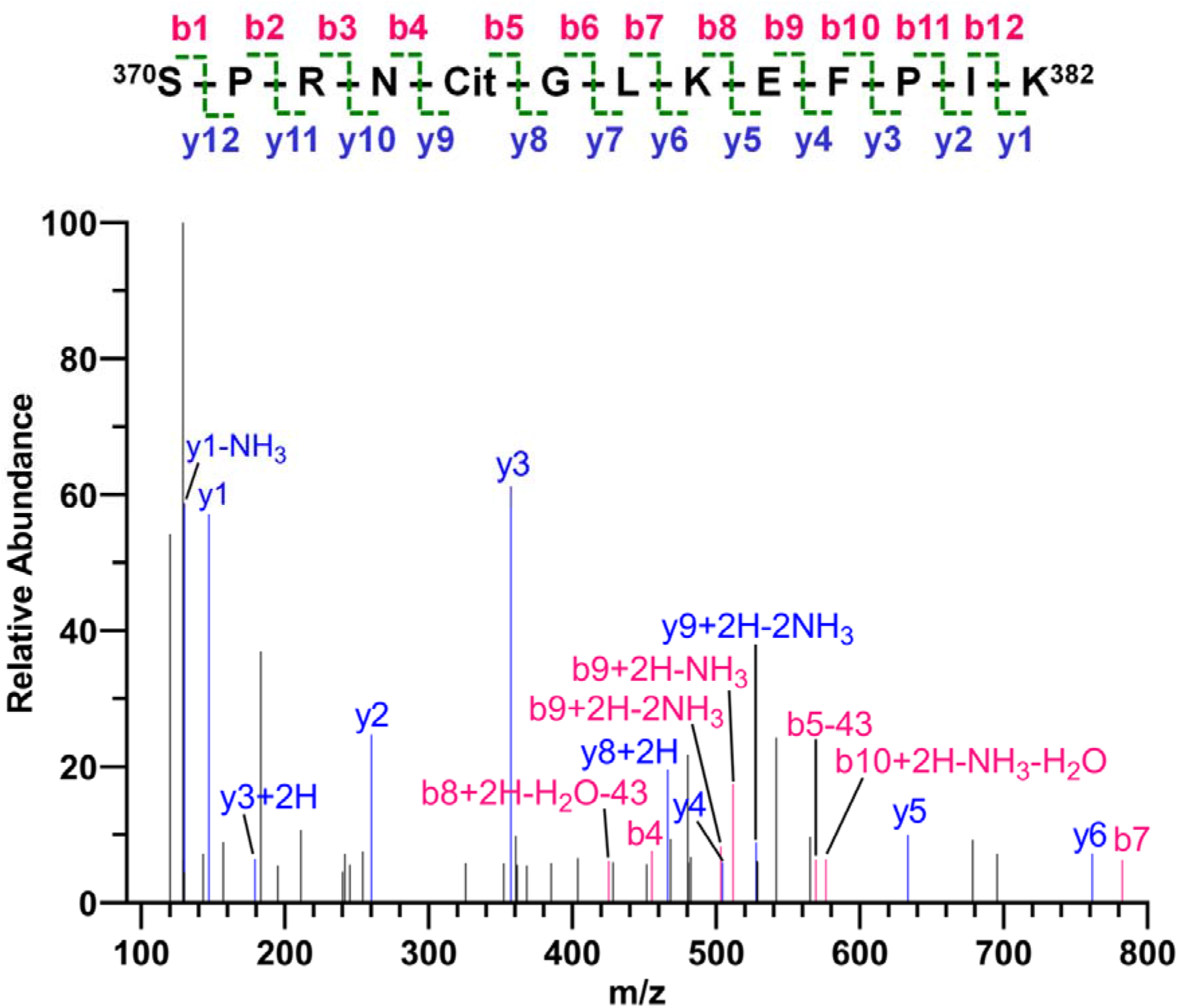
MS2 spectrum of the peptide (^370^SPRN**Cit** GLKEFPIK^382^) containing the Cit374 residue. This peptide was generated by digesting R374Cit mutant PAD4 with Lys-C and Glu-C. A 43 Da neutral loss, characteristic to the citrulline side chain that loses isocyanic acid during fragmentation, from b5, and not from b4 ion confirms citrulline incorporation at the 374 position.

**Figure S12.**
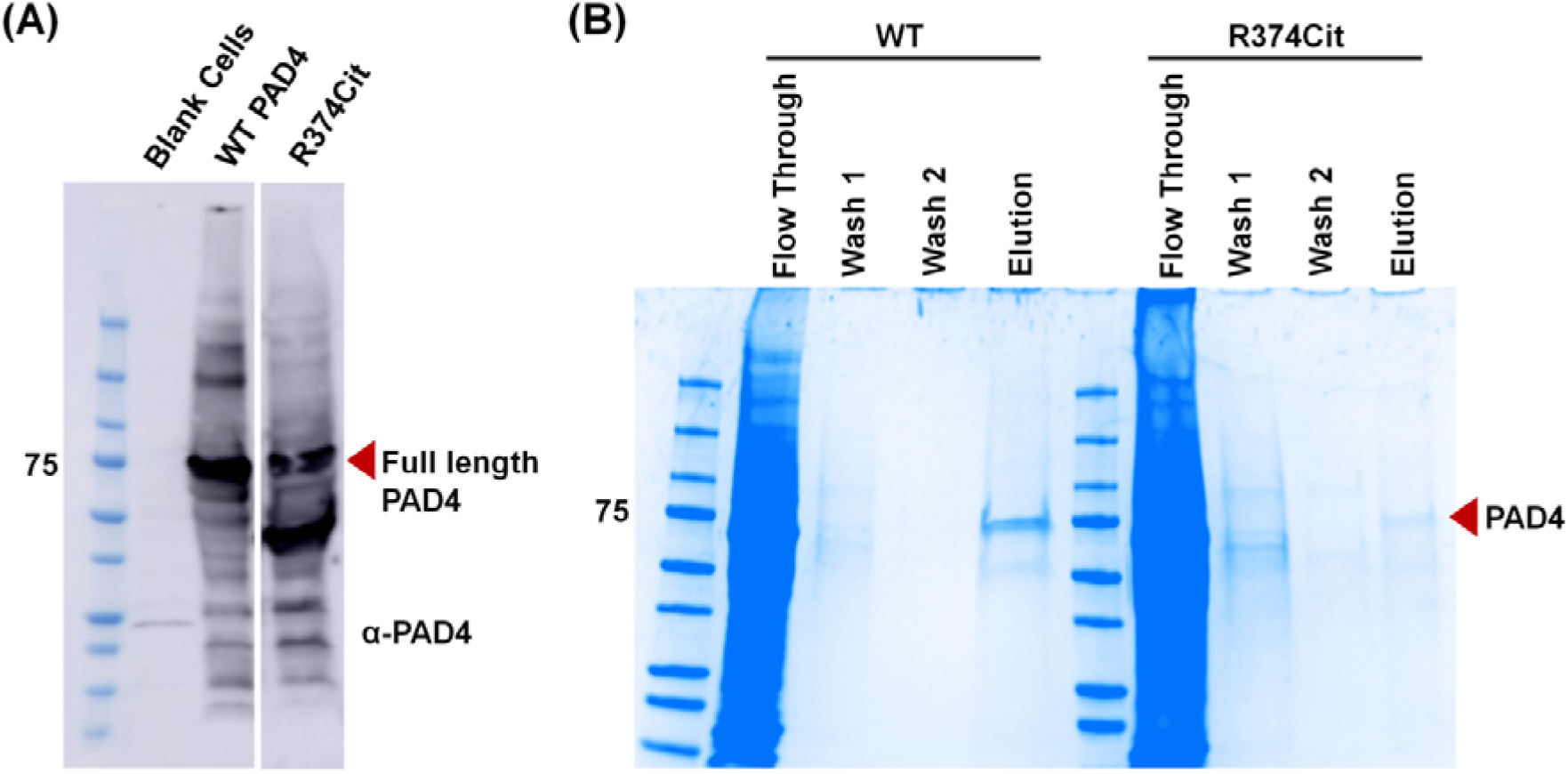
(A) Expression of WT and R374Cit PAD4 in EXPI293F cells as monitored by western blot analysis of the lysate using an α-PAD4 antibody. EXPI293F cell lysate (without transfection) served as a negative control. (B) Coomassie stain of purified WT and R374Cit PAD4 from EXPI293F lysate by Ni-NTA affinity chromatography. Wash 1, Wash 2 and Elution buffer contained 50, 75 and 300 mM imidazole, and the elution fraction was dialyzed to remove the imidazole.

**Figure S13.**
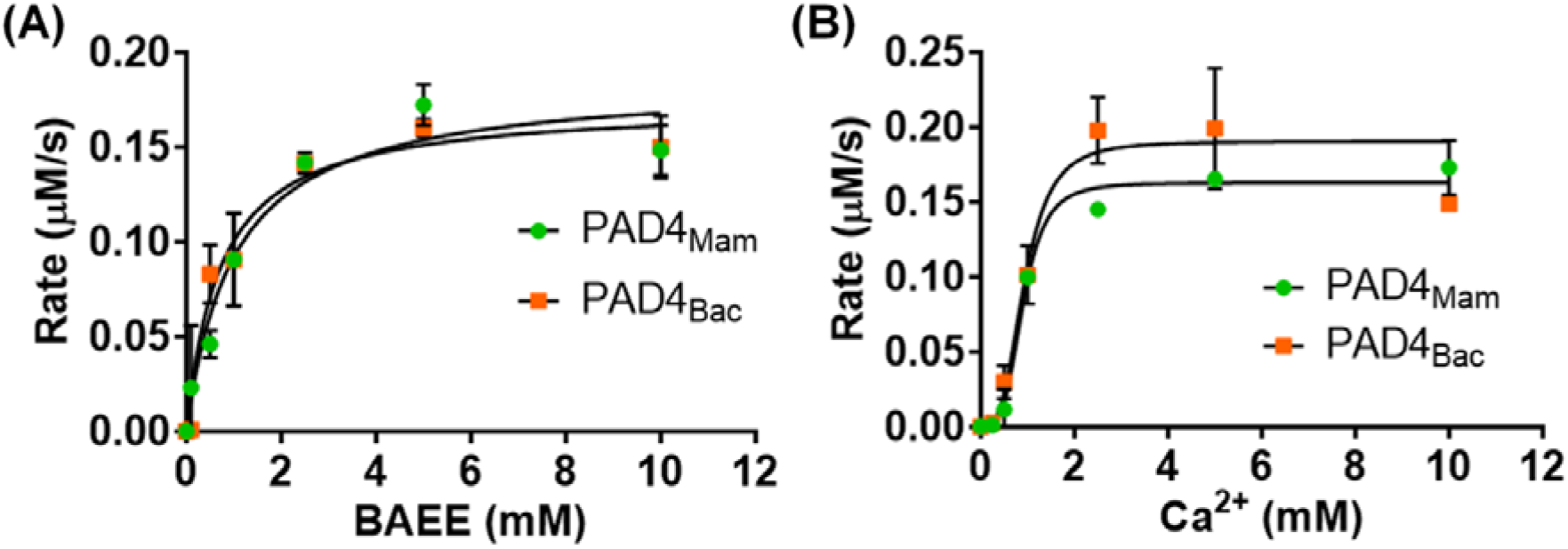
(A) Michaelis-Menten kinetics for the citrullination of BAEE by PAD4_Mam_ (purified from mammalian expression system) and PAD4_Bac_ (purified from bacterial expression system), indicating that both these enzymes have similar steady-state kinetic parameters (Table S3). Since PAD4_Mam_ contains an N-terminal FLAG and a C-terminal His tag, these results also indicate that the presence of these tags do not affect enzymatic activity. (B) Calcium-dependence plots for PAD4_Mam_ and PAD4_Bac_. These data indicate that regardless of the source of the enzyme, the *K*_0.5_ (Ca^2+^ concentration for half-maximal activity) values are 0.9 mM.

**Figure S14.**
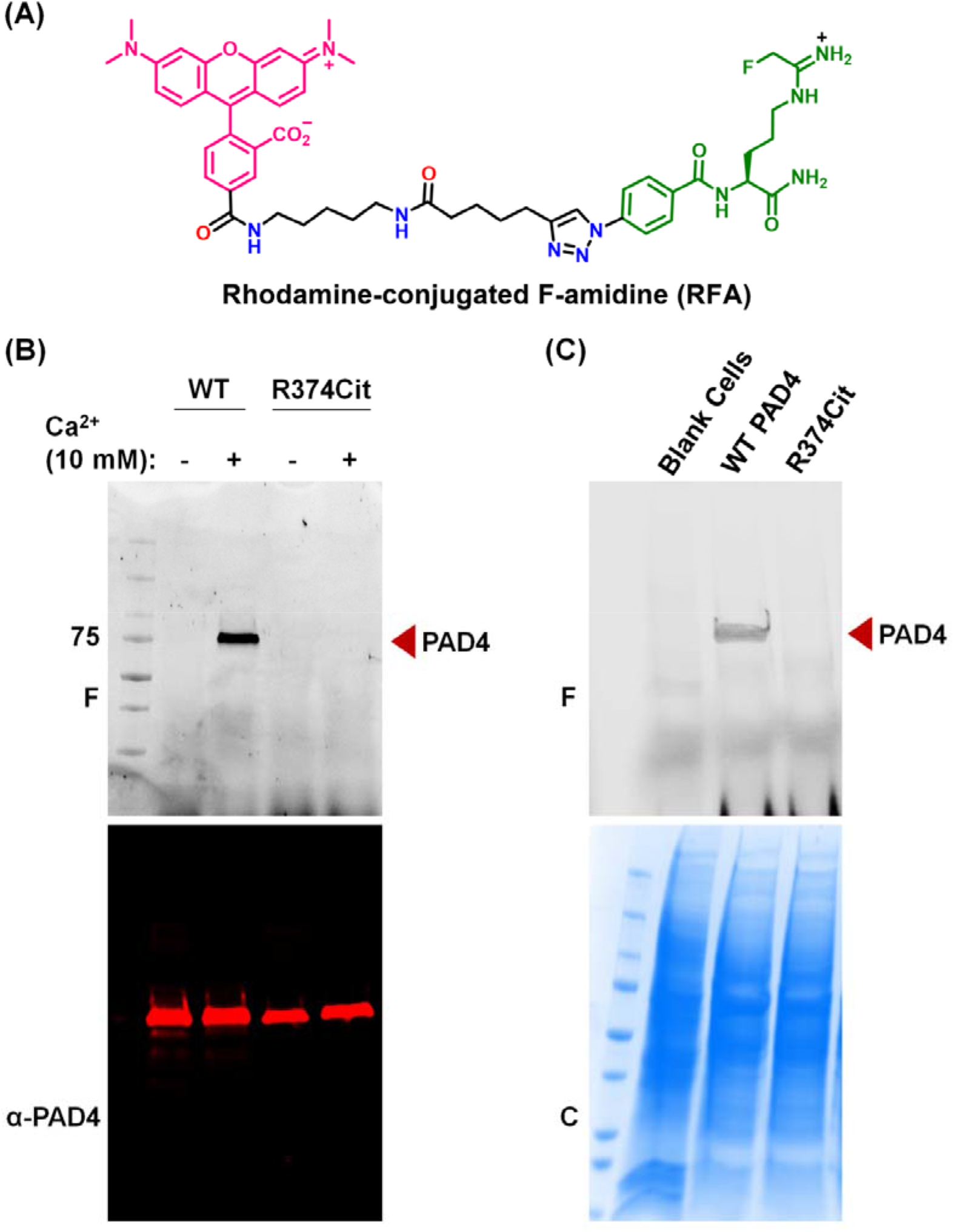
(A) Chemical structure of rhodamine-conjugated F-amidine (RFA). (B) RFA-labeling of wild-type (WT) and R374Cit PAD4 in the presence and absence of Ca^2+^. (C) RFA-labeling of EXPI293F cell lysate containing wild-type (WT), R372Cit and R374Cit PAD4. EXPI293F cell lysate (without transfection) served as a negative control. F and C refer to the fluorograph and coomassie brilliant blue stain, respectively.

**Figure S15.**
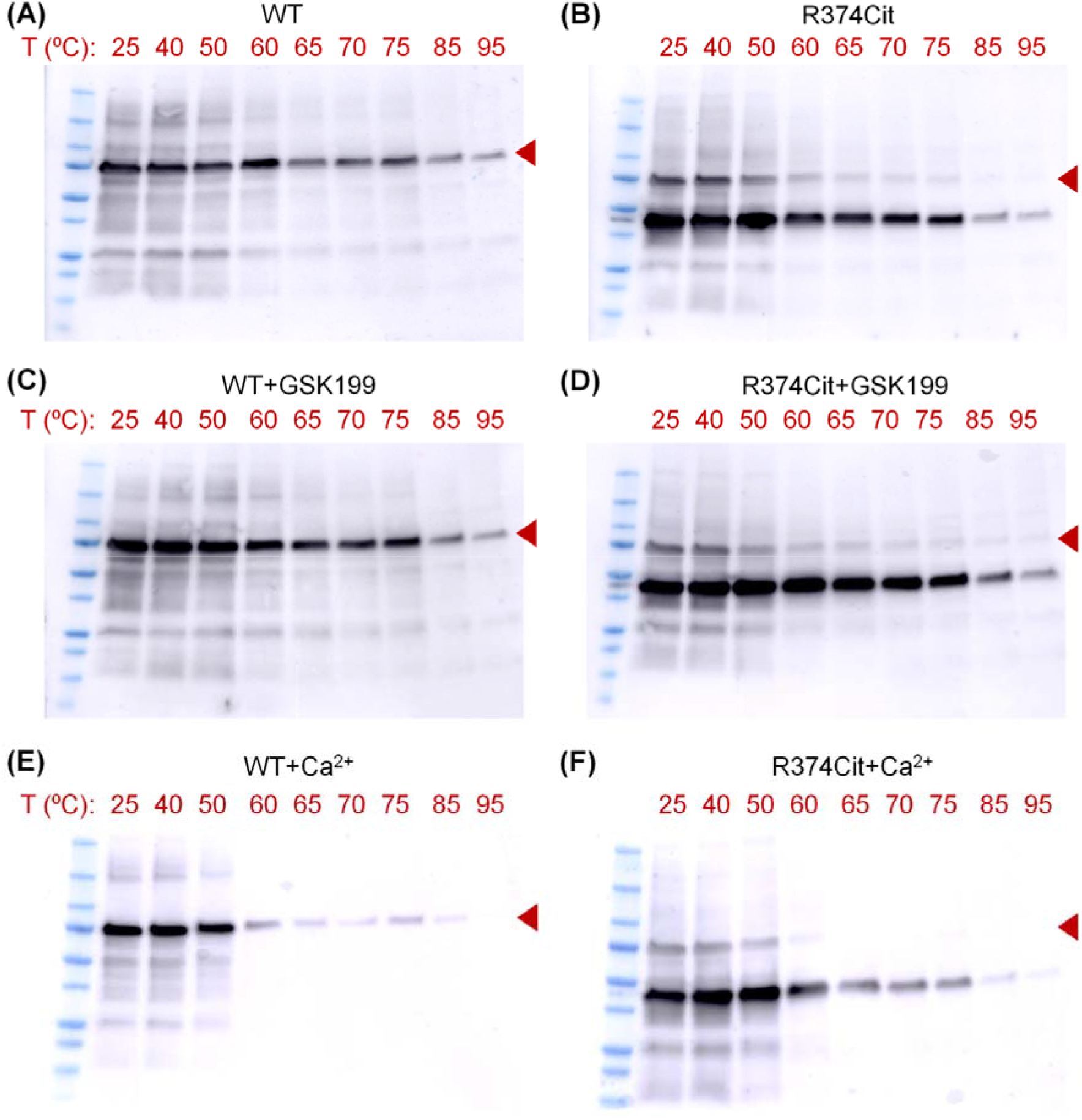
Full western blot images for the thermal shift assays using EXPI293F cell lysate containing wild-type (WT) and R374Cit PAD4. GSK199 and Ca^2+^ were used at 10 μM and 1 mM concentration, respectively. Full-length PAD4 is indicated by a brown triangle in each blot.

**Figure S16.**
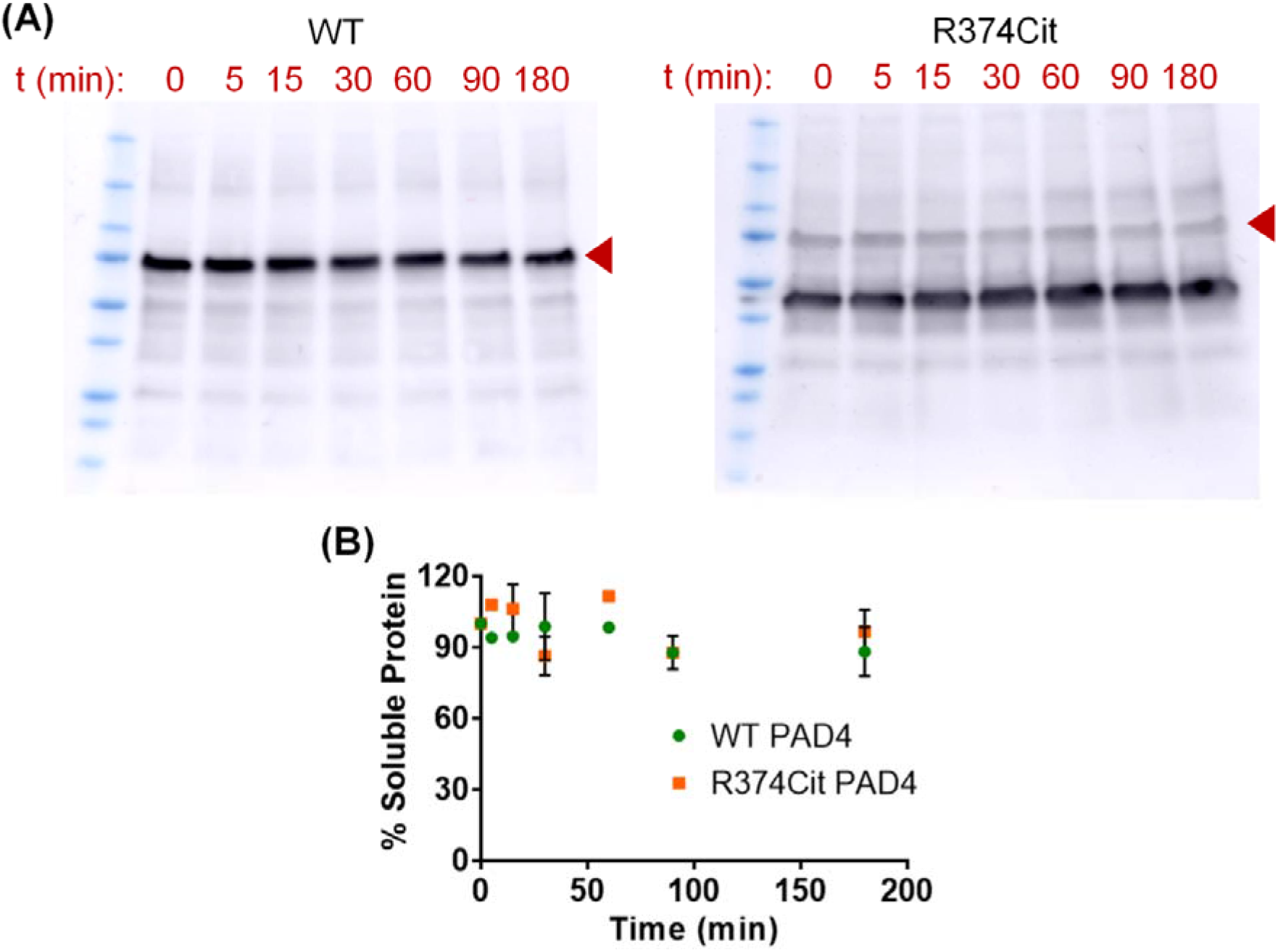
(A) Thermal stability of WT and R374Cit PAD4 at 37 °C over 3 h. EXPI293F cell lysate containing WT PAD4 or R374Cit mutant was used in this assay. Full-length PAD4 is indicated by a brown triangle in each blot. (B) Variation of PAD4 band intensities in panel A over 180 min, indicating that R374Cit mutant is as stable as WT PAD4 at 37 °C.

**Figure S17.**
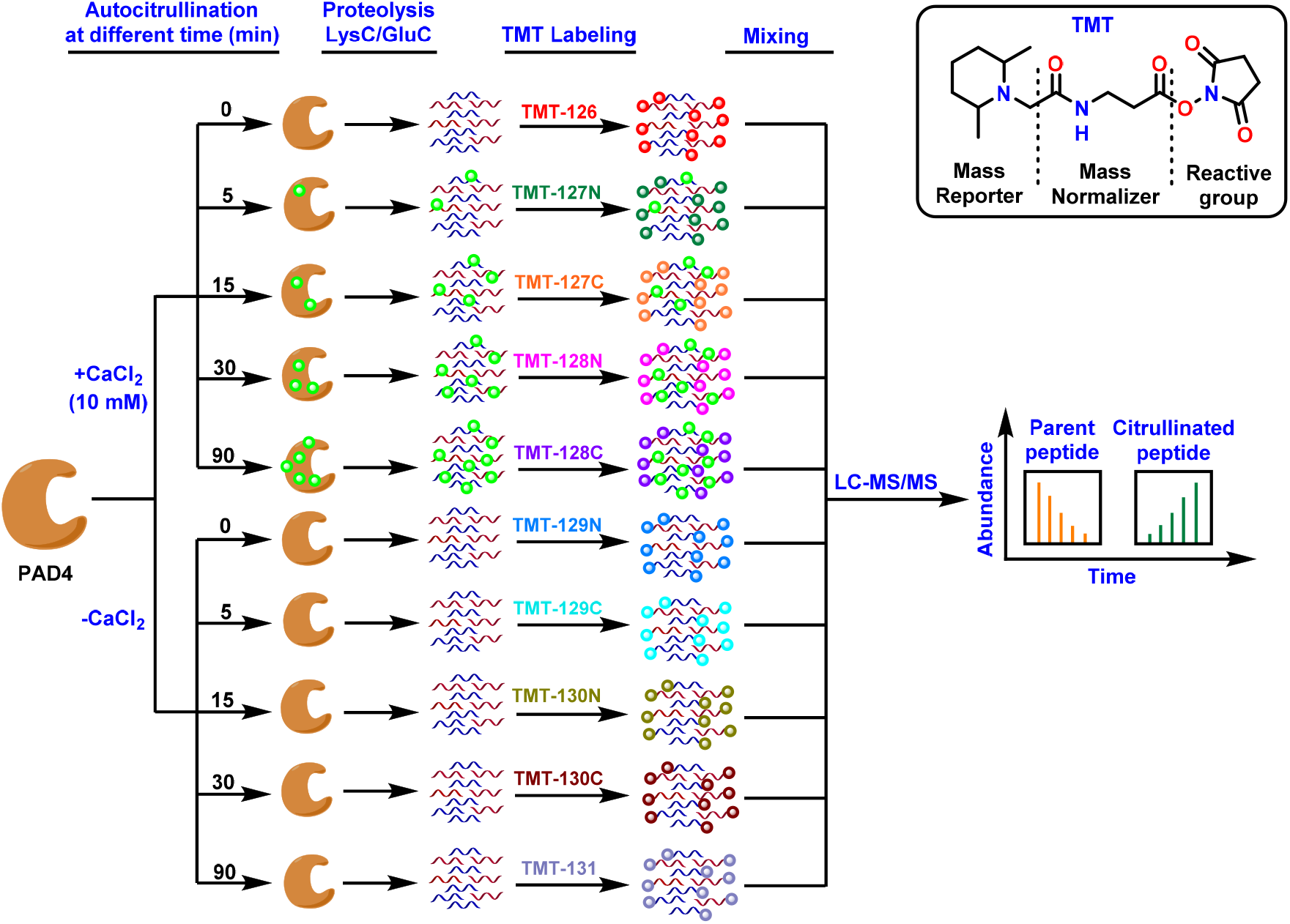
Schematic representation of quantitative proteomic analysis of time-dependent autocitrullination of PAD4. Samples treated in the absence of calcium served as negative controls. Peptides derived from proteolysis of PAD4 with Lys-C and Glu-C were labeled with isobaric tandem mass tags (TMT) to enable simultaneous quantification of autocitrullination at various sites with increasing time.

## Materials and General Methods

N^α^-Fmoc-N^δ^-L-Ornithine hydrochloride, HBTU, HOBt and other Fmoc-protected amino acids were purchased from Chem-Impex International Inc. 1-(isocyanatomenthyl)-2-nitrobenzene (1 M solution in toluene) was purchased from Ellanova Laboratories. Triethylamine, trifluoroacetic acid, anhydrous dichloromethane, anhydrous dimethylformamide, piperidine and HPLC-grade acetonitrile were bought from Sigma-Aldrich. Halt protease inhibitor cocktail (EDTA-free), Universal nuclease, Ni-NTA resin, Pierce™ Peptide Desalting Spin Columns (Catalogue No 89852), Pierce™ Quantitative Fluorometric Peptide Assay kit (Catalogue No 23290) and TMT10plex™ Isobaric Label Reagent (Catalogue No 90110) were obtained from ThermoFisher Scientific. Deuterated solvents were purchased from Cambridge Isotope Laboratories. Plasmid purification kit was bought from Bio Basic Canada Inc. The TOP10 *E. coli* strain was used for plasmid construction and propagation. The cells were grown in LB liquid medium with 100 μg/ml ampicillin or 50 μg/ml kanamycin. Primers were synthesized by Integrated DNA Technologies (Coralville, IA). Restriction enzymes (NEB, Beverly, MA), Phusion Hot Start II DNA polymerase (Fisher Scientific, MA) and T4 DNA ligase (Enzymatics, Beverly, MA) were used for plasmid construction following manufacturers’ protocol. White light and fluorescence imaging of HEK293T cells expressing the EGFP-39-TAG reporter were performed using a Zeiss AX10 microscope. Rabbit polyclonal anti-PAD4 (catalogue no. ab50332) was obtained from Abcam. EXPI293F cells, EXPI293™ expression medium and ExpiFectamine™ 293 Transfection Kit were obtained from Gibco. Dulbecco’s Modified Eagle’s medium (DMEM), fetal bovine serum (FBS) and Antibiotic-Antimycotic (100X) solution were obtained from Gibco and were used for HEK293T cell maintenance. Mass spec grade Lys-C (Catalogue No VA117A) and sequencing grade Glu-C (Catalogue No V165A) was obtained from Promega. ^1^H and ^13^C NMR spectra were recorded in *d_6_*-DMSO as solvent using a Bruker 500 MHz NMR spectrometer. Chemical shift values are cited with respect to SiMe_4_ (TMS) as the internal standard. All the compounds were purified by reverse-phase HPLC using a semi-preparative C18 column (Agilent, 21.2 × 250 mm, 10 μm) and a water/acetonitrile gradient supplemented with 0.05% trifluoroacetic acid. Fluorographs were recorded using a Typhoon scanner with excitation/emission maxima of ∼546/579, respectively. Wild-type PAD4 from bacterial expression system (PAD4_Bac_) were expressed and purified as reported earlier.^3^

### Synthesis

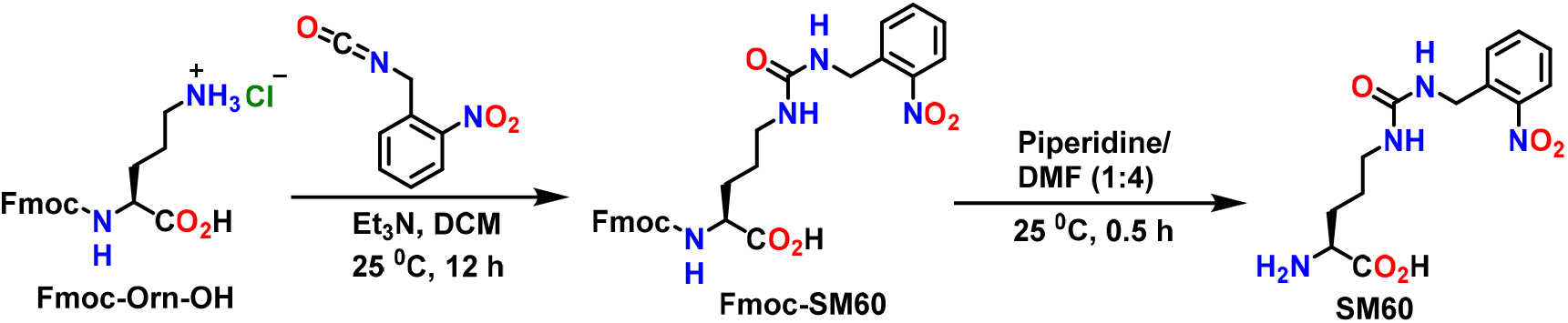
Synthesis of SM60.

Fmoc-Orn-OH (1 g, 2.6 mmol) was suspended in anhydrous dichloromethane and triethylamine (0.7 mL, 5.2 mmol) was added to it. 1-(isocyanatomenthyl)-2-nitrobenzene (1 M solution in toluene) (2.6 mL, 2.6 mmol) was then added dropwise and the reaction mixture was stirred at room temperature for 12 h. Excess triethylamine and dichloromethane was then evaporated under reduced pressure to afford a yellowish brown semisolid that was used for the subsequent step without further purification. The crude product was dissolved in 1:4 piperidine/dimethylformamide (10 mL) and was stirred at room temperature for 30 min. The reaction mixture was then vigorously stirred with excess hexane and the hexane layer was decanted off. Washing with hexane was repeated several times to remove most of the dimethylformamide. The pale yellow semisolid obtained thereafter was purified by reverse phase HPLC using a pre-packed C18 column and a water/acetonitrile (supplemented with 0.05% trifluoroacetic acid) gradient as eluent to afford **SM60** as a white solid (overall yield: 60%). **SM60** was thoroughly characterized with ^1^H and ^13^C NMR spectroscopy and Mass spectrometry. ^1^H NMR (DMSO-*d_6_*) δ (ppm): 8.18 (s, 3H), 7.94 (dd, *J* = 9 Hz, 1H), 7.64-7.67 (m, 1H), 7.43-7.48 (m, 2H), 6.48 (t, *J* = 5 Hz, 1H), 6.24 (t, *J* = 5 Hz, 1H), 4.4 (d, *J* = 10 Hz, 2H), 3.85 (s, 1H), 2.93-2.97 (m, 2H), 1.61-1.75 (m, 2H), 1.40-1.49 (m, 1H), 1.31-1.39 (m, 1H); ^13^C NMR (DMSO-*d_6_*) δ (ppm): 171.5, 158.5, 148.3, 136.6, 134.2, 130.0, 128.4, 124.9, 52.3, 40.7, 39.1, 28.0, 26.2; ESI-MS (m/z) calculated for C_13_H_18_N_4_O_5_ [M + H]^+^: 311.14, found 311.20.

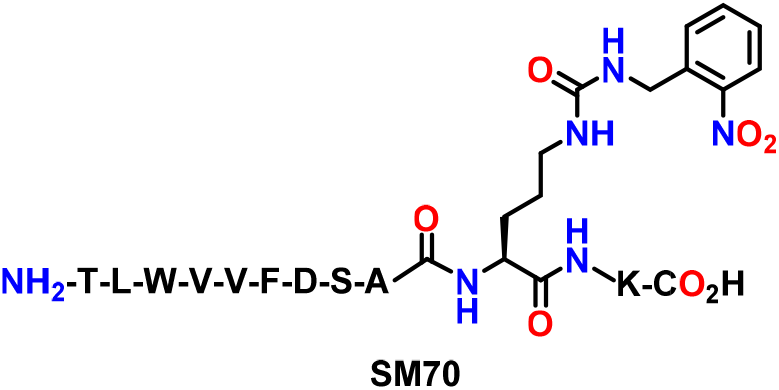
Synthesis of SM70.

**SM70** was synthesized using an automated solid-phase peptide synthesizer (PS3, Protein Technologies, Inc.) by following manufacturer’s protocol. Briefly, Fmoc-Lys(Boc)-Wang resin (350 mg, 0.2 mmol) was taken in a 30 mL glass reaction vessel, and **Fmoc-SM60** (425 mg, 0.8 mmol), Fmoc-Ala-OH (249 mg, 0.8 mmol), Fmoc-Ser(tBu)-OH (306 mg, 0.8 mmol), Fmoc-Asp(OtBu)-OH (329 mg, 0.8 mmol), Fmoc-Phe-OH (310 mg, 0.8 mmol), Fmoc-Val-OH (272 mg, 0.8 mmol), Fmoc-Val-OH (272 mg, 0.8 mmol), Fmoc-Trp(Boc)-OH (421 mg, 0.8 mmol), Fmoc-Leu-OH (283 mg, 0.8 mmol) and Fmoc-Thr-OH (318 mg, 0.8 mmol) were taken in separate amino acid vials. HBTU (303 mg, 0.8 mmol) and HOBt (108 mg, 0.8 mmol) were then added to each amino acid vial. N-methylmorpholine (0.4 M in DMF) was used as a base for the activation of the carboxylic acid group with HBTU and HOBt. The N-terminal Fmoc protecting group on each amino acid was removed with 20% piperidine in DMF. Each peptide coupling reaction was carried out for 1 h. Once all the amino acids were coupled, the resin was transferred to a synthetic column, thoroughly washed with DMF and DCM, and the peptide was cleaved from the resin by treating with a cleavage cocktail (95% trifluoroacetic acid, 4.5% triisopropylsilane, 0.5% water, 10 mL) at room temperature for 30 min with constant mixing. The flow-through was collected and the resin was washed 2-3 times with trifluoroacetic acid. The combined flow through and trifluoroacetic acid washes were then treated with 8-10 times excess cold diethyl ether to precipitate the peptide. Excess ether and trifluoroacetic acid was slowly evaporated by purging nitrogen. The crude peptide was purified by reverse phase HPLC using a pre-packed C18 column and a water/acetonitrile (supplemented with 0.05% trifluoroacetic acid) gradient to afford **SM70** as a white solid (overall yield: 15%). **SM70** was characterized by ESI mass spectrometry, ESI-MS (m/z) calculated for C_69_H_100_N_16_O_19_ [M]^+^: 1456.74, found 1456.60.

### Construction of plasmids

The previously reported pAcBac3-EcLeuTAG-EGFP-39-TAG plasmid was used to construct additional plasmids.^4^ EGFP-39-TAG was replaced with WT PAD4 using SfiI restriction site to create pAcBac3-EcLeuPLRS1TAG-PAD4WT. For incorporation of **SM60**, we introduced TAG nonsense codon at the desired sites by site-directed mutagenesis based on the pAcBac3-EcLeuPLRS1TAG-PAD4WT plasmid. pIDTSMART eRF1 E55D was generated following a paper from the Chin lab.^5^

List of Primers:

PAD4WT-Forward: ATTATTAGAATTGGCCAAGGAGGCCACCATGGACTACAAGGACGACGACGACAA G
PAD4WT-Reverse: ATTATTAGAATTCGGCCTTAGAGGCCTCAGTGGTGGTGGTGGTGGTGGTGGTGGTGGTGGTGGTGGGGCACCATGTTCCACCACTTGAA
PAD4 R372TAG Inner Forward: GTCTTCGACTCTCCTTAGAACTAGGGCCTGAAG
PAD4 R372TAG Inner Reverse: CTTCAGGCCCTAGTTCTAAGGAGAGTCGAAGAC
PAD4 R374TAG Inner Forward: GTCTTCGACTCTCCTAGGAACTAGGGCCTGAAG
PAD4 R374TAG Inner Reverse: CTTCAGGCCCTAGTTCCTAGGAGAGTCGAAGAC

### HEK293T cell culture and transfection

HEK293T cells were maintained in 37 °C humidified incubator supplemented with 5 % CO_2_. Cells were seeded as 9 × 10^6^ cells per 10 cm plate 24 h before transfection. EGFP and WT PAD4 transfections were performed by incubating 10 μg plasmid DNA, 50 μL PEI (1 mg/mL; Polysciences, Warrington, PA), and 180 μL DMEM for 10 min at room temperature, followed by adding the solution dropwise to the culture medium of the cells. For **SM60** incorporation into PAD4 at 372 and 374 positions, 12 μg of PAD4 R372TAG or PAD4 R374TAG and 8 μg of pIDTSMART eRF1 E55D plasmids were incubated with a mixture of 100 μL PEI and 180 μL DMEM for 10 min at room temperature before adding to cells. **SM60** was added at the same time to a final concentration of 1 mM, and 2 mM sodium butyrate was added to enhance protein expression.

### EGFP fluorescence analysis

EGFP fluorescence was analyzed 48 h after transfection. DMEM was exchanged with PBS and the plates were irradiated at 365 nm (120 Watt, 10 cm × 10 cm LED array; Larson Electronics) for 75 s at 4 °C to decage **SM60**. Cells were then harvested and resuspended in 600 μL CelLytic M buffer (Sigma, St. Louis, MO) with Halt Protease inhibitor Cocktail (Thermo Scientific, Waltham, MA) and Pierce Universal Nuclease for Cell Lysis (Fisher Scientific, Hampton, NH). Lysate were clarified by centrifugation at 16,000 xg for 10 min and 100 μL of supernatant was transferred to a clear-bottom 96-well plate for fluorescence measurement following previously described protocol.^4^ All the experiments were performed at least in duplicate.

### Overexpression of wild-type and mutant (R374Cit) PAD4 in EXPI293F Cells

EXPI293F cells were cotransfected with the engineered LeuRS/tRNALeu/PAD4 and eRF genes using ExpiFectamine™ 293 transfection kit by following the manufacturer’s protocol. Briefly, EXPI293F cells were grown to 2.9 × 10^6^ density in the EXPI293™ expression medium (9 mL) at 37 °C under 5% CO_2_ atmosphere. pAcBac3 Plasmid (6 μg) encoding the genetically engineered LeuRS, tRNA^Leu^ and WT PAD4 or mutant PAD4 (containing TAG mutation at 374 position) and the plasmid encoding release factor, i.e. eRF (4 μg), were resuspended in 0.5 mL opti-MEM™ I reduced serum media. ExpiFectamine™ (27 μL) was also resuspended in 0.5 mL opti-MEM™ media and then the plasmid mixture was slowly added to ExpiFectamine™ solution. This mixture was incubated at room temperature for 20 min to from the DNA polyplex. Then the DNA polyplex (1 mL) was slowly transferred to the EXPI293F cell suspension (9 mL) and **SM60** (100 μL of 100 mM stock in DMSO, 1 mM final) was added to the medium. The cells were then incubated at 37 °C under 5% CO_2_ atmosphere for 24 h with constant shaking. Transfection enhancers 1 (50 μL) and 2 (500 μL) (supplied with the ExpiFectamine™ 293 transfection kit) were then added to the cell suspension and the cells were further grown at 37 °C under 5% CO_2_ atmosphere for 48 h with constant shaking. Cells were then harvested, washed with cold Dulbecco's Phosphate-Buffered Saline (DPBS), resuspended in 5 mL DPBS and were taken in a cell culture dish (100 mm × 20 mm, Corning). Cells were then irradiated with 365 nm light for 5 minute using a photoreactor (Luzchem) containing 14 UV-A lamps (8 W each). After UV-A irradiation, cells were harvested, resuspended in 1 mL in DPBS (containing 1X Halt protease inhibitor cocktail) and lysed using a probe sonicator. Overexpression of wild-type and mutant PAD4 was confirmed by western blot analysis of the EXPI293F cell lysate using rabbit polyclonal anti-PAD4 antibody.

### Purification of wild-type and mutant PAD4 from HEK293T cells

Cells from a 10 cm plate were harvested 48 h after transfection. For cells that overexpressed proteins containing **SM60**, media was exchanged with PBS and the plates were irradiated at 365 nm (120 Watt,10 cm × 10 cm LED array (Larson Electronics)) for 75 s at 4 °C to decage **SM60** right before harvesting. The cells were resuspended in 600 μL CelLytic M buffer (Sigma, St. Louis, MO) with Halt protease inhibitor cocktail (Thermo Scientific, Waltham, MA) and Pierce universal nuclease for cell lysis (Fisher Scientific, Hampton, NH). After a 10 min incubation at room temperature, 1.2 mL of equilibration buffer (20 mM Na_2_HPO_4_, 300 mM NaCl, 10 mM imidazole pH 7.4) was added. Lysate was clarified by centrifugation at 16,000 g for 10 min at 4 °C. The clarified cell-free extract was subjected to Ni-NTA affinity chromatography using HisPur resin (Fisher Scientific, Hampton, NH) following the manufacturer’s protocol.

### Purification of wild-type and mutant PAD4 from EXPI293F cells

Following UV-A irradiation, cells were resuspended in lysis buffer (20 mM TRIS-HCl pH 8.0, 400 mM NaCl, 10 mM imidazole, 2 mM DTT, 10 % glycerol, 1X protease inhibitor cocktail, 1X universal nuclease) and lysed using a probe sonicator. The lysate was centrifuged at 18000 × *g* for 20 min at 4 °C and the supernatant was incubated with Ni-NTA agarose beads (prewashed with the lysis buffer) for 30 min at 4 °C on an end-over-end shaker. The beads were then transferred to a synthetic column and washed sequentially with buffer 1 (20 mM TRIS-HCl pH 8.0, 400 mM NaCl, 50 mM imidazole, 2 mM DTT, 10 % glycerol) and buffer 2 (20 mM TRIS-HCl, pH 8.0, 400 mM NaCl, 75 mM imidazole, 2 mM DTT, 10 % glycerol). Finally PAD4 was eluted from the beads using elution buffer (20 mM TRIS-HCl pH 8.0, 500 mM NaCl, 300 mM imidazole, 2 mM DTT, 10 % glycerol). Eluted PAD4 was then dialyzed against 20 mM TRIS-HCl, pH 8.0, 500 mM NaCl, 2 mM DTT, 10 % glycerol using a dialysis cassette with a molecular weight cut off 3.5 kDa.

### Digestion of R374Cit PAD4 and LC-MS/MS Analysis

15 μg of R374Cit mutant was resuspended in 20 mM TRIS-HCl (200 μL, pH 7.4) and trichloroacetic acid (20% final) was added to it. The cloudy mixture was vortexed vigorously and was kept at −20 °C for 30 min to precipitate the protein. Then the mixture was centrifuged at 15,000 rpm for 30 min at 4 °C. The pellet was washed with cold acetone, resuspended in 30 μL of 8 M urea in PBS (pH 7.4) with sonication and 70 μL of 100 mM ammonium bicarbonate was added to the solution. Then 1.5 μL of 1 M DTT was added and the solution was incubated at 65 °C for 20 min. 2.5 μL of freshly-prepared 500 mM iodoacetamide was added and the solution was incubated at room temperature for 30 min in dark. The alkylation reaction was quenched by 120 μL PBS (pH 7.4). 1 μL Lys-C (1 μg/μL, reconstituted in water) and 2 μL Glu-C (0.5 μg/μL, reconstituted in water) was then added to the solution and the mixture was incubated at 37 °C for 16 h. The proteolysis was terminated by adding 10 μL of formic acid. The peptide mixture was then desalted using Pierce™ desalting C18 spin columns following manufacturer’s protocol and was analysed by LC-MS/MS as described later.

### Time-dependent citrulline production by wild-type, R372Cit and R374Cit PAD4 purified from mammalian expression system

PAD4 (6 μL of 1 μM stock, 100 nM final) was added to a pre-warmed (10 min, 37 °C) reaction mixture (60 μL final) containing N^α^-benzoyl arginine ethyl ester (BAEE, 10 mM), CaCl_2_ (10 mM), TRIS-HCl (100 mM, pH 7.4), NaCl (50 mM) and DTT (2 mM). This mixture was incubated at 37 °C for 0, 30, 45 and 90 min after which the reaction was stopped by flash freezing with liquid nitrogen. The production of citrulline at various time points was quantitated by COLDER assay.^6,7^ The time-dependent citrulline production by PAD4 was fit to the equation of a straight line using Graphpad Prism. All the reactions were performed at least in duplicate.

### Michaelis-Menten kinetics of wild-type and mutant PAD4

PAD4 (50 nM final for wild-type, 75 nM final for R372Cit and 100 nM final for R374Cit) was added to a pre-warmed (10 min, 37 °C) reaction mixture (60 μL final) containing various concentrations of BAEE (0, 0.5, 1, 2.5, 5 and 10 mM), CaCl_2_ (10 mM), TRIS-HCl (100 mM, pH 7.4), NaCl (50 mM) and DTT (2 mM). This mixture was incubated at 37 °C for 45 min (for wild-type), 90 min (for R374Cit) and 120 min (for R372Cit) followed by quenching with liquid nitrogen. The rate of citrulline formation at various BAEE concentrations was quantitated with the COLDER assay.^6,7^ The rates were plotted against the BAEE concentration and were fit to the Michaelis-Menten equation using Graphpad Prism. All the reactions were performed at least in duplicate.

### Calcium-dependence of wild-type PAD4

PAD4_Bac_ (purified from bacterial expression system) and PAD4_Mam_ (purified from mammalian expression system) (6 μL of 0.5 μM stock, 50 nM final) were added to separate pre-warmed (10 min, 37 °C) reaction mixture (60 μL final) containing BAEE (10 mM), various concentrations of CaCl_2_ (0, 0.25, 0.5, 1, 2.5, 5 and 10 mM), TRIS-HCl (100 mM, pH 7.4), NaCl (50 mM) and DTT (2 mM). This mixture was incubated at 37 °C for 45 min followed by flash freezing with liquid nitrogen. The production of citrulline at various concentration of CaCl_2_ by PAD4 was quantified with the COLDER assay.^6,7^ The calcium-dependence of citrulline production by PAD4 was fit to the equation 1,

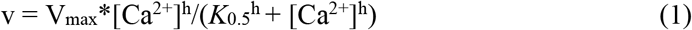

using Graphpad Prism, where v is the velocity of reaction, V_max_ is the maximum velocity of reaction, [Ca^2+^] is the concentration of calcium, h is the hill slope and *K*_0.5_ is the calcium concentration that gives half-maximal velocity. All the reactions were performed at least in duplicate.

### Effect of autocitrullination of wild-type PAD4 on the enzymatic activity

PAD4_Bac_ (0.5 μM final) was added to a pre-warmed (10 min, 37 °C) solution containing CaCl_2_ (0 or 10 mM), TRIS-HCl (100 mM, pH 7.4), NaCl (500 mM), DTT (2 mM). At various time points (0, 5, 15, 30, 60 and 90 min), 6 μL of this reaction mixture was removed and was added to a pre-warmed (10 min, 37 °C) reaction mixture (10 mM BAEE, 10 mM CaCl_2_, 100 mM TRIS, pH 7.4, 500 mM NaCl, 2 mM DTT, with a final volume of 60 μL). After 45 min, the reaction mixture was flash frozen with liquid nitrogen. The production of citrulline at various time points was quantified with the COLDER assay.^6,7^ The loss in activity of PAD4 over time was fit into single exponential decay using Graphpad Prism. All the reactions were performed at least in duplicate.

### Rhodamine-PG labeling of autocitrullinated PAD4

Rhodamine-PG labeling of autocitrullinated PAD4 was performed as reported earlier with minor modifications.^8^ PAD4 (0.5 μM final) was added to a pre-warmed (10 min, 37 °C) reaction mixture containing CaCl_2_ (0 or 10 mM), TRIS-HCl (100 mM, pH 7.4), NaCl (500 mM), DTT (2 mM). At various time points (0, 5, 15, 30, 60 and 90 min), the reaction was stopped by flash freezing in liquid nitrogen. The samples were thawed, and trichloroacetic acid (20% final) and rhodamine-PG (80 μM final) were sequentially added to it. The reaction mixture was incubated at 37 °C for 1 h. Then the reaction was quenched with citrulline (100 mM final) dissolved in 50 mM TRIS-HCl (pH 7.4) and the mixture was further incubated at 37 °C for 30 min. The samples were placed at −20 °C for 30 min and the precipitated proteins were collected by centrifugation (15000 rpm for 30 min) at 4 °C. The protein pellet was washed with cold acetone and dried. Then the pellet was dissolved in 40 μL buffer containing 100 mM arginine, 20 mM TRIS-HCl (pH 7.4), 1% SDS and 10 μL 5X SDS loading dye. The proteins were separated by SDS-PAGE and the fluorescently labeled bands were visualized by scanning the gel in a typhoon scanner (excitation and emission maxima ~546 and 579 nm, respectively). The fluorescent intensities of protein bands were quantified using ImageJ software and were normalized against the coomassie intensities that indicate the amount of protein present in each lane. All the reactions were performed at least in duplicate.

### RFA labeling of wild-type and R374Cit PAD4

RFA labeling of PAD4 was carried out by following a protocol similar to that established for PAD1 and PAD2.^9,10^ Briefly, PAD4 (100 nM final) was added to pre-warmed (10 min, 37 °C) reaction mixture (100 mM TRIS pH 7.4, 500 mM NaCl, 0 or 10 mM CaCl_2_, and 2 mM DTT in a final volume of 30 μL) containing RFA (200 nM final). After incubating at 37 °C for 2 h, the reaction mixture was quenched with 5X SDS-PAGE loading dye and boiled at 95 °C for 10 min. The proteins were separated by SDS-PAGE using a 4-20% gradient gel and fluorescently labeled proteins were visualized by scanning the gel in a typhoon scanner (excitation and emission maxima ~546 and 579 nm, respectively). The fluorescent intensities of protein bands were quantified using ImageJ software. All the reactions were performed at least in duplicate.

### RFA labeling of wild-type and R374Cit PAD4 in EXPI293F lysate

RFA (10 μM) was added to a pre-warmed (10 min, 37 °C) reaction mixture (2 mg/mL EXPI293F lysate containing wild-type or R374Cit or R372Cit PAD4 in 1X PBS, 2 mM CaCl_2_, and 2 mM DTT in a final volume of 50 μL) and the mixture was incubated at 37 °C for 2 h. The reaction was quenched with 5X SDS-PAGE loading dye and was boiled at 95 °C for 10 min. The proteins were separated on a 4-20% SDS-PAGE gel and the fluorescently labeled proteins were visualized by scanning the gel in a typhoon scanner (excitation and emission maxima ~546 and 579 nm, respectively). The fluorescent intensities of protein bands were quantified using ImageJ software. All the reactions were performed at least in duplicate.

### Thermal shift Assay

A 50 μL reaction mixture (EXPI293F lysate containing overexpressed wild-type PAD4 (1.5 mg/mL) or R374Cit mutant (2 mg/mL), 1X PBS and DMSO (1% final)) was heated at various temperatures (25, 40, 50, 60, 65, 70, 75, 85 and 90 °C) for 5 min followed by flash freezing with liquid nitrogen. The samples were thawed and the precipitated proteins were separated by centrifugation (15000 rpm, 30 min) at 4 °C. 40 μL of the supernatant was mixed with 10 μL of 5X SDS-PAGE loading dye and the mixture was boiled at 95 °C for 10 min. The proteins were separated by SDS-PAGE using a 4-20% gradient gel and the soluble fractions of PAD4 were quantitated by western blot analysis using a rabbit polyclonal anti-PAD4 antibody. This assay was also performed separately in the presence of CaCl_2_ (1 mM final) and a PAD4-selective ligand, GSK199 (10 μM final). For these assays, the reaction mixture was incubated with CaCl_2_ or GSK199 at room temperature for 5 min before heating up at different temperatures. All the reactions were performed at least in duplicate.

### Proteomics study on autocitrullinated PAD4

50 μg PAD4 was incubated at 37 °C for various times (0, 5, 15, 30 and 90 min) in the absence and presence (10 mM) of CaCl_2_ followed by flash freezing with liquid nitrogen. The samples were thawed and trichloroacetic acid (20% final) was added. Then the samples were placed at −20 °C for 30 min and the precipitated proteins were collected by centrifugation (15000 rpm, 30 min) at 4 °C, washed with cold acetone and dried. The protein pellet was resuspended in 6 M urea (100 μL) in PBS and TCEP (2 mM final) was added to it. The solution was incubated at 37 °C for 1 h. Then iodoacetamide (4 mM final) was added and the solution was incubated at 37 °C for 30 min in dark. Then 200 μL of PBS was added to the solution to achieve a final concentration of urea of 2 M. 1 μL Lys-C (1 μg/μL, reconstituted in water) and 2 μL Glu-C (0.5 μg/μL, reconstituted in water) was then added to the samples and the samples were incubated at 37 °C for 16 h. The proteolysis was terminated by adding 15 μL of formic acid. The peptide mixture was then desalted with Pierce™ desalting C18 spin columns, resuspended in 120 μL HEPES buffer (100 mM, pH 8.5) and the total peptide content in each sample was quantified and normalized by the Pierce™ Quantitative Fluorometric Peptide Assay kit by following manufacturer’s protocol. 8 μL of 19.5 μg/μL acetonitrile stock of TMT10plex™ isobaric labeling reagents (TMT10-126, TMT10-127N, TMT10-127C, TMT10-128N, TMT10-128C, TMT10-129N, TMT10-129C, TMT10-130N, TMT10-130C and TMT10-131) were added to 100 μL samples treated in the absence and presence of calcium for 5 different time points (0, 5, 15, 30 and 90 min). The reaction mixtures were incubated at room temperature for 1 h. To quench the reaction, 8 μL of 5% hydroxylamine was added to each sample and the mixture was incubated at room temperature for 15 min. Then the 10 samples (0, 5, 15, 30 and 90 min samples in the absence and presence of calcium) were combined together in a new microcentrifuge tube and desalted using C18 spin columns. This experiment was performed in triplicate.

### LC-MS/MS analysis

Peptides were lyophilized, re-suspended in 5% acetonitrile, 0.1% (v/v) formic acid in water, and loaded at 4.0 μL/min by a NanoAcquity UPLC (Waters Corporation, Milford, MA) onto a 100 μm I.D. fused-silica pre-column packed with 2 cm of 5 μm (200 Å) Magic C18AQ (Bruker-Michrom), equilibrated with 5% acetonitrile, 0.1% (v/v) formic acid in water. After trapping for 4.0 minutes on the pre-column, peptides were eluted at 300 nL/min from a 75 μm I.D. gravity-pulled analytical column packed with 25 cm of 3 μm (100Å) Magic C18AQ particles using a gradient of mobile phase A, 0.1% (v/v) formic acid in water and mobile phase B, 0.1% (v/v) formic acid in acetonitrile as follows; 0-100 min (5-35 %B), 100-120 min (35-65 %B), 120-121 min (65-95 %B), and 121-126 min (95 %B). Ions were introduced by positive electrospray ionization via liquid junction at 1.4 kV into a Q Exactive hybrid quadrapole orbitrap mass spectrometer (Thermo Scientific, Waltham, MA). Mass spectra were acquired over *m/z* 300-1750 at 70,000 resolution (*m/z* 200) with an AGC target of 1e6, and data-dependent acquisition selected the top 10 most abundant precursor ions for tandem mass spectrometry by HCD fragmentation using an isolation width of 1.6 Da, max fill time of 100 ms, and AGC target of 1e5. Peptides were fragmented by a normalized collisional energy (NCE) of 27 and product ion spectra acquired at a resolution of 17500 (*m/z* 200). For TMT-labeled samples, NCE was set to 32 and product ion spectra were acquired at a resolution of 35000 (*m/z* 200).

### LC-MS/MS Data analysis

Raw data files were peak processed with Proteome Discoverer (version 2.1, ThermoScientific, Waltham, MA) followed by identification using Mascot Server (version 2.5, Matrix Science) against the Swissprot human or E.coli (TMT-labeled samples) FASTA file. Proteolytic enzyme was set to LysC and GluC with two missed cleavages. Variable modifications of N-terminal acetylation, oxidized methionine, pyroglutamic acid for glutamine, deamidation of asparagine, and citrullination of arginine were implemented. Carbamidomethylation of cysteines and TMT6-plex modification at lysine and peptide N-terminus were set as fixed modifications. Assignments were made using a 10 ppm mass tolerance for the precursor and 0.05 Da mass tolerance for the fragments. All non-filtered search results were processed by Scaffold (version 4.10.0, Proteome Software, Inc.) utilizing the Trans-Proteomic Pipeline (Institute for Systems Biology) with threshold values set at 90% for peptides (1% false-discovery rate) and 99% for proteins (2 peptides minimum, 6% false-discovery rate). TMT product ion ratios were calculated using the Scaffold Q+S analysis software.

